# Systematic creation and phenotyping of Mendelian disease models in *C. elegans*: towards large-scale drug repurposing

**DOI:** 10.1101/2023.08.25.554786

**Authors:** Thomas J. O’Brien, Ida L. Barlow, Luigi Feriani, André E.X. Brown

## Abstract

There are thousands of Mendelian diseases with more being discovered weekly and the majority have no approved treatments. To address this need, we require scalable approaches that are relatively inexpensive compared to traditional drug development. In the absence of a validated drug target, phenotypic screening in model organisms provides a route for identifying candidate treatments. Success requires a screenable phenotype, however the right phenotype and assay may not be obvious for pleiotropic neuromuscular disorders. Here we show that high-throughput imaging and quantitative phenotyping can be conducted systematically on a panel of *C. elegans* disease model strains. We used CRISPR genome-editing to create 25 worm models of human Mendelian diseases and phenotyped them using a single standardised assay. All but two strains were significantly different from wild-type controls in at least one feature. The observed phenotypes were diverse, but mutations of genes predicted to have related functions in their human orthologs led to similar behavioural differences in worms. As a proof-of-concept, we performed a drug repurposing screen of an FDA approved compound library, and identified two compounds that rescued the behavioural phenotype of a model of UNC80 deficiency. Our results show that a single assay to measure multiple phenotypes can be applied systematically to diverse Mendelian disease models. The relatively short time and low cost associated with creating and phenotyping multiple strains suggests that high-throughput worm tracking could provide a scalable approach to drug repurposing commensurate with the number of Mendelian diseases.

## Introduction

By definition, rare genetic diseases each affect a small number of patients. However, because there are many kinds of rare diseases, they are collectively common and affect more than 300 million people worldwide (Nguengang Wakap et al., 2020). Over 80% of rare genetic diseases affect children and many have severe symptoms (Bavisetty et al., 2013). Furthermore, 95% of rare diseases have no approved treatment (Nguengang Wakap et al., 2020). Given the severity and prevalence of rare diseases, the lack of treatments represents an important unmet need. Genome and exome sequencing has become faster and cheaper, enabling its widespread use in the diagnosis of genetic disease. In turn, this has led to the rapid identification of genetic lesions associated with rare genetic diseases (Gonzaga-Jauregui et al., 2012). Genetic diagnosis is an important first step, but often presents patients and physicians with a research problem rather than a treatment.

Genetic diagnoses rarely lead directly to a drug target hypothesis, especially for loss-of-function mutations. In the absence of a validated drug target, phenotypic screens in a model organism provide an alternative route to candidate treatments. Three things are required for a phenotypic screen to be successful: (1) a conserved gene; (2) a genetically tractable organism or culture system, in which the disease mutation can be introduced; and (3) a measurable phenotype compatible with high-throughput screening. For the model nematode *C. elegans*, more than half of the human genes associated with rare genetic diseases in the OMIM database are conserved (Kropp et al., 2021) and CRISPR-based editing has made the generation of *in vivo* models containing disease-relevant mutations faster and easier. However, the third condition, a screenable phenotype, remains a bottleneck for many mutations and models.

If all three conditions are met, a phenotypic screen for candidate treatments can be performed. Because there is limited investment in drug development for rare diseases, lead optimisation and safety testing may not be possible. Hence, there is a particular focus on screening approved drugs that have already been shown to be safe and bioavailable in humans to reduce the cost and time it takes to translate screening hits to human trials (Roessler et al., 2021). Two recent examples of promising repurposing screens using multiple model organisms illustrate this potential.

Firstly, modelling amyotrophic lateral sclerosis (ALS) with disease-associated genetic variants of conserved genes in *C. elegans*, zebrafish and mice led to the discovery and clinical trials for repurposing of the antipsychotic drug, pimozide. By exploiting high throughput drug screening in *C. elegans* carrying point mutations in *TDP-43*, a panel of drugs were identified that rescued ALS movement and neurodevelopment phenotypes in all animal models. Early clinical trials found pimozide improved clinical and physiological outcomes in ALS patients (Patten et al., 2017). Secondly, the glycosylation disorder, PMM2-CDG, is caused by mutations in *PMM2* and presents clinically with developmental delay, psychomotor retardation and axial hypotonia, among other symptoms. *C. elegans* carrying the same disease-associated mutation in their endogenous *pmm-2* gene exhibited larval arrest upon pharmacological ER stress. *In vivo* screening identified two compounds that rescued development in *C. elegans* and enzymatic activity in PMM2-CDG patient fibroblasts (Iyer et al., 2019). One of the compounds, the aldose reductase inhibitor epalrestat, is currently being tested in clinical trials for PMM2-CDG patients (National Clinical Trial number: NCT04925960).

A screenable phenotype was critical for the success of these studies. In the ALS model, worms became paralysed after 2 hours in liquid and in the PMM2-CDG model, brood size was severely affected. The majority (∼74%) of rare genetic diseases affect nervous system function (Lee et al., 2020), and many of these mutations will not cause strong effects on development or gross motility when mutated in worms. In this study, we use a combination of high-throughput imaging (Barlow et al., 2022) and quantitative phenotyping (Javer, Currie, et al., 2018; Javer, Ripoll-Sánchez, et al., 2018) to phenotype a diverse panel of 25 *C. elegans* strains with mutations in orthologs of human disease genes. All but two of the mutant strains had detectable phenotypes compared to controls in at least one aspect of their morphology, posture, or motion. No single feature was different in all strains highlighting the need for a multidimensional phenotype. Mutation of genes predicted to have similar functions in their human orthologs (e.g., *bbs-1*, *bbs-2,* and *tub-1*) led to similar phenotypes. As a proof-of-concept, we then performed a repurposing screen using a library of 743 FDA approved compounds to identify drugs that improved the behavioural phenotype of *unc-80* loss-of-function mutants. Liranaftate and atorvastatin rescued the core behavioural features associated with *unc-80* loss of function and did not cause a large number of detectable side effects.

The ability to detect phenotypic difference in diverse strains using a standardised 16-minute assay will make it possible to perform repurposing screens for existing and newly described rare diseases efficiently.

## Results

### Selection and molecular characterisation of a diverse panel of C. elegans disease models

The problem of large-scale disease modelling can be thought of as a problem of sampling and connecting two related genotype-phenotype (GP) maps (Fig. 1A). The first is the human disease map. The genomic space is sampled by mutation and mating in humans and the map is elucidated by sequencing and clinical phenotyping of patients with genetic diseases. The second is the model organism map. Here, the genomic space is sampled using mutagenesis and genome editing and the map is elucidated using laboratory experiments. Given the accelerated characterisation of the human GP map, we sought to accelerate the characterisation of the corresponding *C. elegans* GP map. We follow a phenolog-inspired approach to connecting the two maps. Provided there is a causal connection between a genetic variant and a disease in humans and that the causal gene is conserved in worms, the observed worm phenotype may be a useful disease model even if the connection to the human phenotype is non-obvious (McGary et al., 2010).

**Figure 1:**
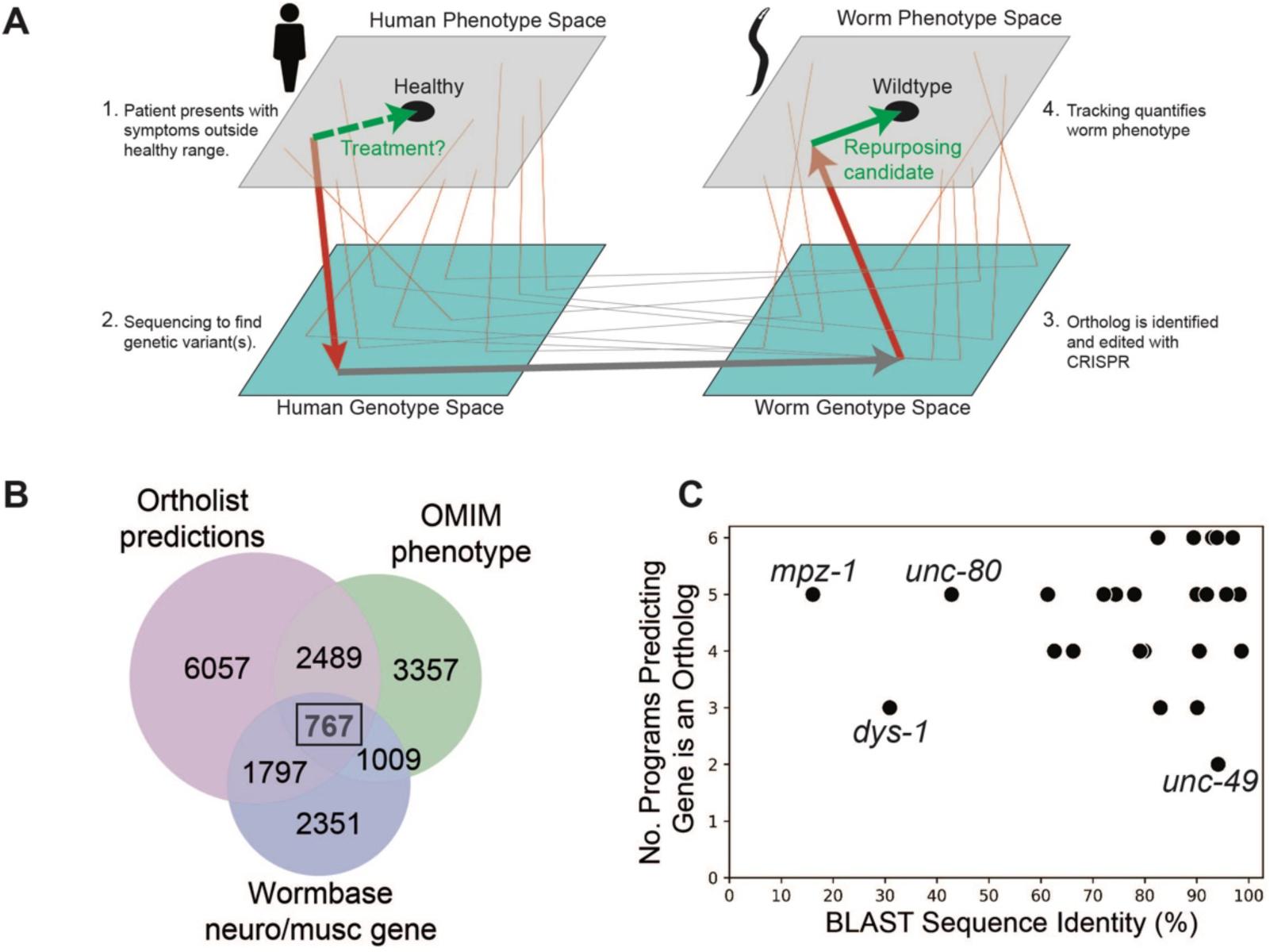
Overview of genotype-phenotype mapping and disease model panel. Disease modelling as genotype-phenotype mapping in humans and a model organism. Arrows show progression from symptom identification (a phenotype outside the healthy range) to genotyping (placing a patient in genotype space by identifying a genetic variant), disease model creation (making a corresponding mutation in a model organism), and model organism phenotyping. Phenotypic drug screens identify compounds that move a disease model towards the wild-type phenotype (green arrow). These candidates can then be tested in humans (dashed green arrow). Thin lines show other symptom-gene-ortholog-phenotype connections. **(B)** Venn diagram showing number of conserved genes (Ortholist 2), those involved in neuron or muscle function (Wormbase), and those associated with human genetic disorders according to the Online Mendelian Inheritance in Man (OMIM) database. **(C)** Sequence similarity between human and C. elegans genes and the total number of orthology programs predicting that the gene is an ortholog.

To identify a set of *C. elegans* genes that samples a broad genotypic space with connections to the human disease GP map, we first filtered the full database of human-*C. elegans* gene orthologs, Ortholist 2 (Kim et al., 2018), using three criteria: (1) at least two orthology prediction algorithms agree the human and worm genes are orthologs; (2) the WormBase (version WS270) (Harris et al., 2020) gene description includes either ‘neuro’ or ‘musc’ (this captures variants of neuronal, neural, muscle, muscular etc.); and (3) mutations in the human gene are associated with a Mendelian disease. A total of 767 *C. elegans* genes with 2558 orthologous human protein-coding genes met these criteria (Figure 1B). We then searched the list of high-confidence disease-associated orthologs using the following keyword filters to obtain 543 *C. elegans* genes well suited for the study of human Mendelian neuro-muscular diseases: neuron; neuronal; muscle; muscular; neurotransmission; ion; ion channel; dopamine; serotonin; glutamate; acetylcholine; 5-HT; behaviour; behaviour; disease; epilepsy; autism; Parkinson; schizophrenia; bipolar; ADHD; seizure; G protein coupled receptor; GPCR; coupled receptor; antidepressant; antipsychotic. In the present study, we prioritised genes associated with autosomal recessive disease in humans to increase the chance that a gene deletion would yield an appropriate model, and we focussed on those where loss-of-function mutants were known or likely to be viable. To get the list of 25 genes presented in this study, the final selection criterion was subjective interestingness based on our prior knowledge, WormBase gene descriptions, and/or brief literature searches.

In the final panel of 25 worm genes selected, 22 genes have >60% sequence similarity to their human ortholog and 11 of the genes share >90% sequence similarity to their human counterpart (Figure 1C). Furthermore, 24/25 genes are predicted to be orthologous to human genes across >3 orthology prediction algorithms (Figure 1C), and all selected genes have a BLAST E-value (McGinnis & Madden, 2004) smaller than 3×10^-12^ (Supplementary Table 1).

Large CRISPR-Cas9 deletions (mean 4.4 kb) were made in each of the target genes to generate 25 strains containing a >55% deletion (average 76%) of the chosen gene. We direct the reader to the strain-specific gene cards (Supplementary Information) for specific details on the size and position of genomic deletions for each of the individual mutants as well as a brief summary of the associated disease and worm phenotypes. Although we have focused on genes expected to have direct effects on neurons or muscles, even this small subset of disease-associated genes is predicted to affect diverse cellular processes including neurotransmission, excitability, development, and cellular structure (Table 1). According to OMIM, the mutated genes are associated with 31 rare genetic disorders in humans including intellectual disability, developmental delay, and disorders affecting the muscular and/or nervous systems, and are associated with >70 clinical presentations of disease according to the Human Disease Ontology database (Schriml et al., 2022) (Table 1).

**Table 1.**
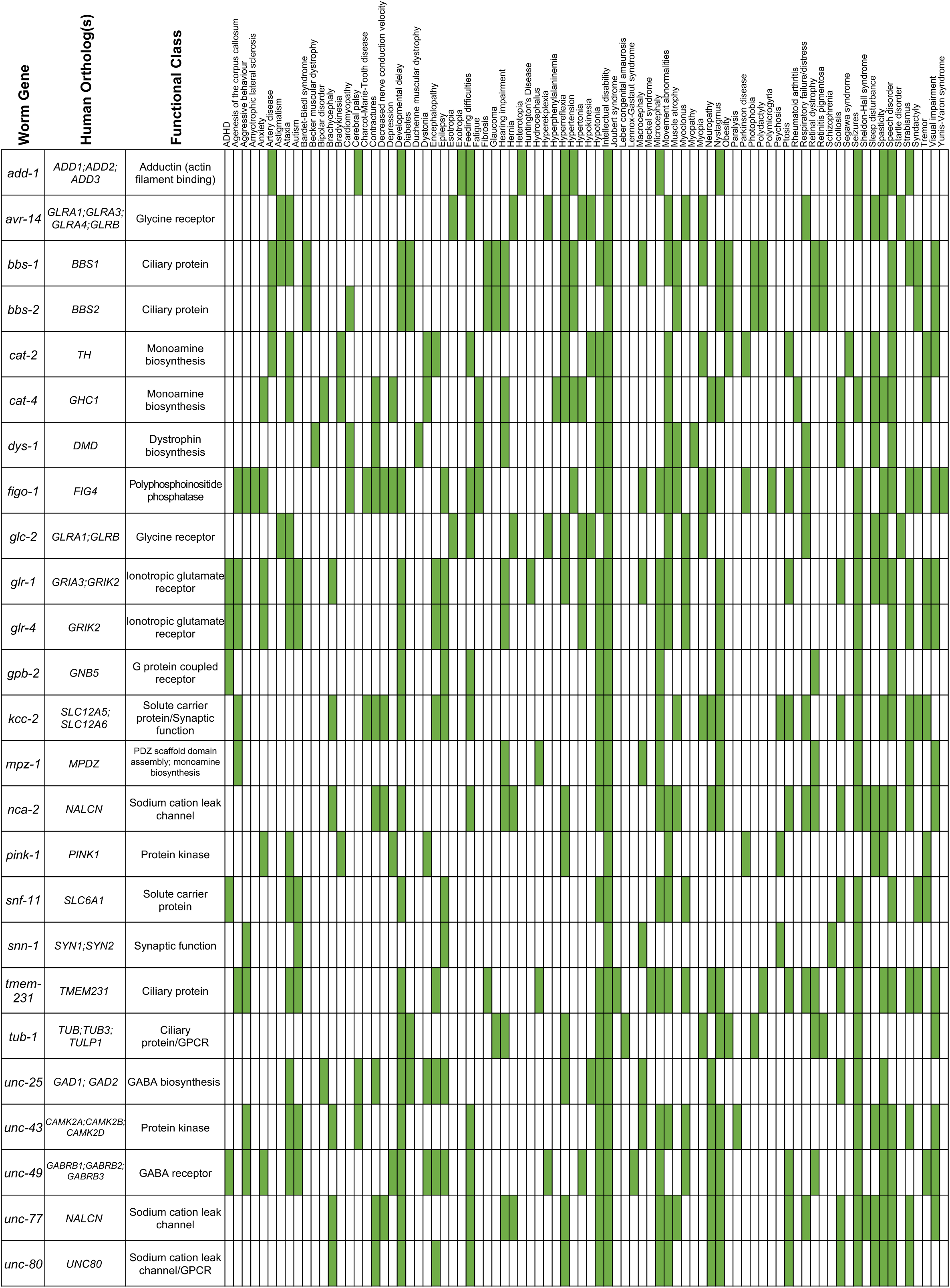
Summary of the associated human ortholog(s), predicted functional class and key associated human disease phenotypes (Human Phenotype Ontology database) for each C. elegans disease model mutant.

### Disease-associated mutations result in diverse phenotypes that are captured by multidimensional behavioural fingerprints

Given that there are thousands of rare diseases, methods to characterise the corresponding disease models should be as scalable as possible. Scalability is improved if a single standardised assay can be used to detect model phenotypes regardless of the mutation. High-resolution worm tracking and behavioural fingerprinting is a promising candidate for a widely applicable assay because it is both fast and multidimensional: a single minutes-long recording can capture differences in morphology, posture, and behaviour that can classify the effect of mutations and drug treatments (Baek et al., 2002; Barlow et al., 2022; Geng et al., 2004; Javer, Ripoll-Sánchez, et al., 2018; McDermott-Rouse et al., 2021; Perni et al., 2018; Ramot et al., 2008; Restif et al., 2014; Swierczek et al., 2011; Tsibidis & Tavernarakis, 2007; Wang & Wang, 2013; Yemini et al., 2013).

We tracked disease model worms on 96-well plates (Figure 2A) in three periods: (1) a 5-minute baseline recording (prestim); (2) a 6-minute video with three ten-second blue light stimuli separated by 90 seconds (blue light), and (3) a 5-minute post-stimulus recording (poststim). For each recording we extract 2763 features covering morphology, posture, and motion and concatenate the feature vectors together to represent the phenotype of each strain (8289 features in total). Many of these features are highly correlated and so for clustering we use a pre-selected subset of 256 features (Javer, Ripoll-Sánchez, et al., 2018) (768 across the three recordings). Using these reduced feature vectors, we performed hierarchical clustering across all strains which demonstrates phenotypic diversity (many of the strains are both different from the N2 wild-type strain and from each other) as well as some similarities. For example, *bbs-1, bbs-2* and *tub-1* mutants affect cilia formation and result in similar phenotypes that cluster together. The same is true for *cat-2*, *kcc*-2, *snf-11* and *unc-25* mutants, which all affect neurotransmitter biosynthesis, GABA transport or the regulation of GABAergic synaptic transmission. Two interactive heatmaps, one containing all behavioural feature vectors extracted for every disease model strain and one showing the Tierpsy256 feature set shown in Figure 2C, are also available for download (doi: zenodo.org/records/13911801).

**Figure 2:**
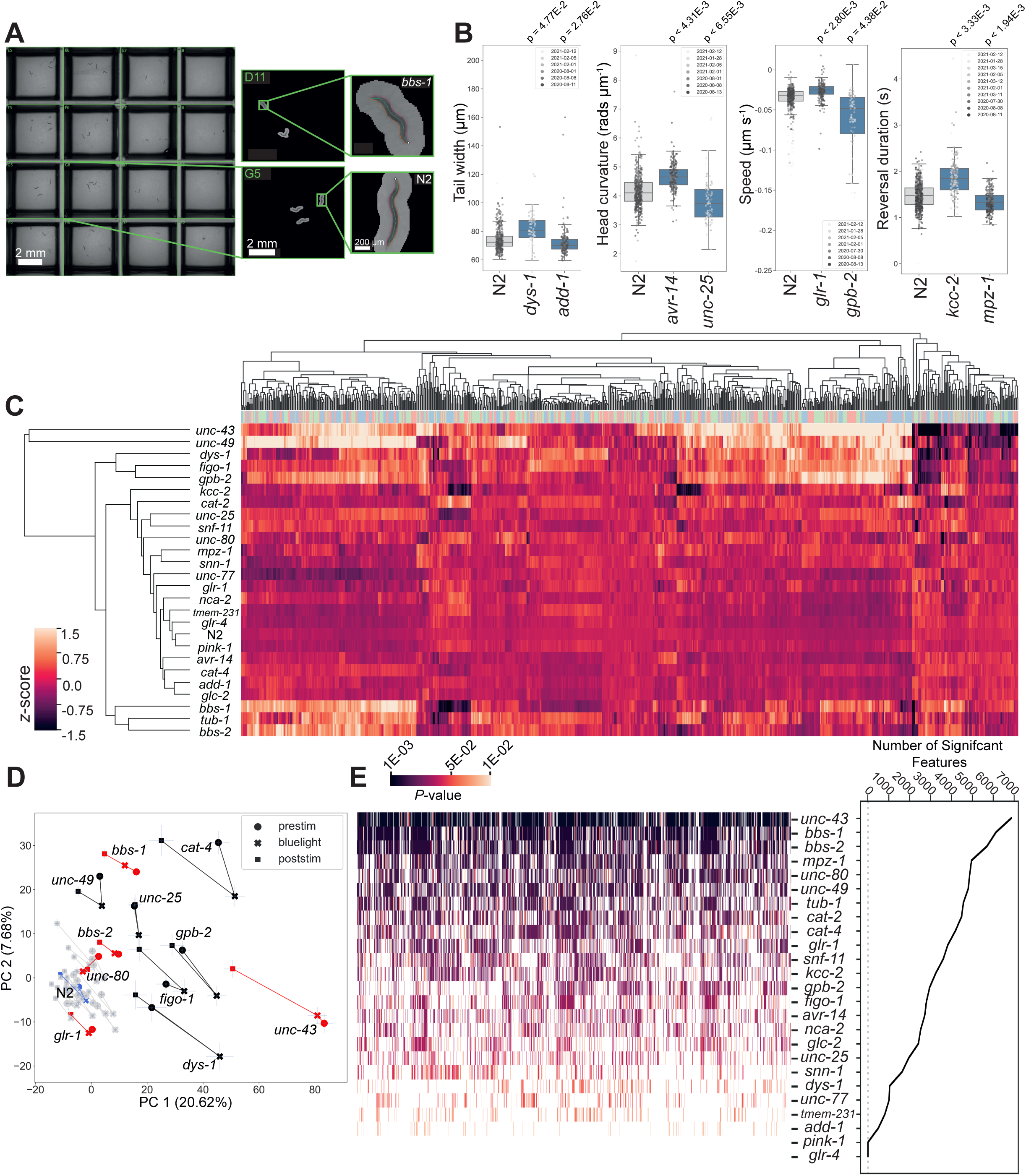
Diverse multidimensional behavioural phenotypes are obtained across the diverse panel of disease model mutants. **(A)** Representative field of view for a single camera channel with individual skeletonised worm images of *bbs-1(syb1588)* (top) and N2 (bottom) imaged on the same plate viewed using Tierpsy Tracker. **(B)** Representative behavioural phenotypes extracted by Tierpsy, representing changes in morphology, posture and locomotion of different disease mutant strains. *P*-values are for comparisons to wild-type N2 worms, block permutation t-test, corrected for multiple comparisons. **(C)** Hierarchical clustering of behavioural fingerprints. Features are *Z*-normalised. The top barcode shows the period of image acquisition where the behavioural feature was extracted by Tierpsy: pre-stimulation (pink), blue light (blue) and post-stimulation (green). **(D)** Principal component analysis of the disease model mutants and N2 reference (blue). Strains move in phenospace between pre-stimulus (circular points), blue light (crosses) and post-stimulation (squares) recordings. Strains with an aberrant blue light response are shown in red. Error bars represent standard error of the mean. **(E)** The total number of statistically significant behavioural features for each strain compared to N2 (from the total set of 8289 features extracted by Tierpsy). *P*-values for each feature were calculated using block permutation t-tests, using n = 100,000 permutations and *P* < 0.05 considered statistically significant after correcting for multiple comparisons using the Benjamini-Yekutieli method.

Principal component analysis (PCA) of the same data, but now separated into prestimulus, blue light, and poststimlus recordings again shows the separation of many of the mutant strains from the wild-type N2 (Figure 2D). The effect of the blue light stimulus on behaviour is clear from the motion of the points through this phenotype space and also reveals new phenotypes. While most strains show a partial recovery in the post-stimulus recording, some strains including *bbs-1*, *bbs-2*, *glr-1*, and *unc-43* display a sustained photophobic response (Figure 2D).

From the panel of 25 disease model strains, 23 are significantly different from wild-type in at least one feature (Figure 2E). The strains that are most different from wild-type show significant differences across a large number of features. These strains are particularly useful for high-throughput drug screens because the phenotypes can be reliably detected with a small number of replicates. Strain-specific summaries (gene cards) of each disease model mutant, its associated phenotype, and individual molecular characteristics are available in the supplementary information. Here we provide more detailed characterisation of two classes of mutants modelling ciliopathies and channelopathies in humans.

### Ciliopathies

Heritable mutations that affect cilia function are associated with a group of rare genetic disorders (ciliopathies) that share diverse symptoms including retinal dystrophy, developmental polydactyly, obesity, cognitive impairment, and renal dysfunction (Reiter & Leroux, 2017). The pleotropic nature of ciliopathies is highlighted by significant interfamilial and intrafamilial phenotypic variability (Shaheen et al., 2016), and the resulting poor gene-phenotype correlation significantly complicates the discovery of effective therapeutics. As a result, current treatment regimens are based on treating individual symptoms.

In the longlist of 543 disease genes, there were 35 genes (6.4% of the total list) predicted to be involved in cilia function. We selected four genes (*bbs-1*, *bbs-*2, *tmem-231* and *tub-1*) involved in regulating nonmotile (primary) cilia development and function that have confirmed expression in the cilia of amphid sensory neurons (Taylor et al., 2021; Ward et al., 2008; Zhang et al., 2022).

Of the genes affecting cilia function, two are orthologs of the Bardet-Biedl syndrome (BBS) family of genes (*bbs-1* and *bbs-2*). BBS is a pleiotropic syndrome often diagnosed in late childhood and is characterised by retinitis pigmentosa, cognitive impairment, obesity, renal dysfunction, hypogonadism, and polydactyly (Forsythe & Beales, 2013). The most prevalent BBS mutations are in *BBS1*, accounting for 23.4% of cases (Forsyth & Gunay-Aygun, 2023), and with 50% embryonic lethality and highly variable phenotypes, *Bbs1-/-* mouse models demonstrate model validity but have not yet yielded insights into potential therapies (Forsyth & Gunay-Aygun, 2023). Mutations in *BBS2* account for 8% of BBS cases (Forsythe & Beales, 2013) and are also implicated in other ciliopathies, such as Meckel syndrome (Karmous-Benailly et al., 2005).

Mutations in *bbs-1* and *bbs-2* led to strong phenotypes including changes in morphology, posture, and locomotion (Figure 3A-D). *bbs-1(syb1588)* and *bbs-2(syb1547)* mutants were shorter, wider and had decreased curvature compared to wild-type worms (Figures 3A, 3B and 3C). They are also hyperactive, with a greater fraction of worms moving during pre-stimulus (baseline) recordings (see individual gene cards), and had faster body bends (derivative of curvature) compared to N2 (Figure 3D and 3F). BBS proteins also regulate the *C. elegans* photoreceptor protein LITE1 in ASH photosensory neurons through a DLK-MAPK signalling pathway independent of their function in cilia (Zhang et al., 2022). Consistent with this, we find that *bbs-1(syb1588)* and *bbs-2(syb1547)* mutants have attenuated blue light sensitivity, with both strains exhibiting a delayed forward, but enhance backward photophobic escape response upon 10-second stimulation with blue light (Figures 3F-H).

**Figure 3:**
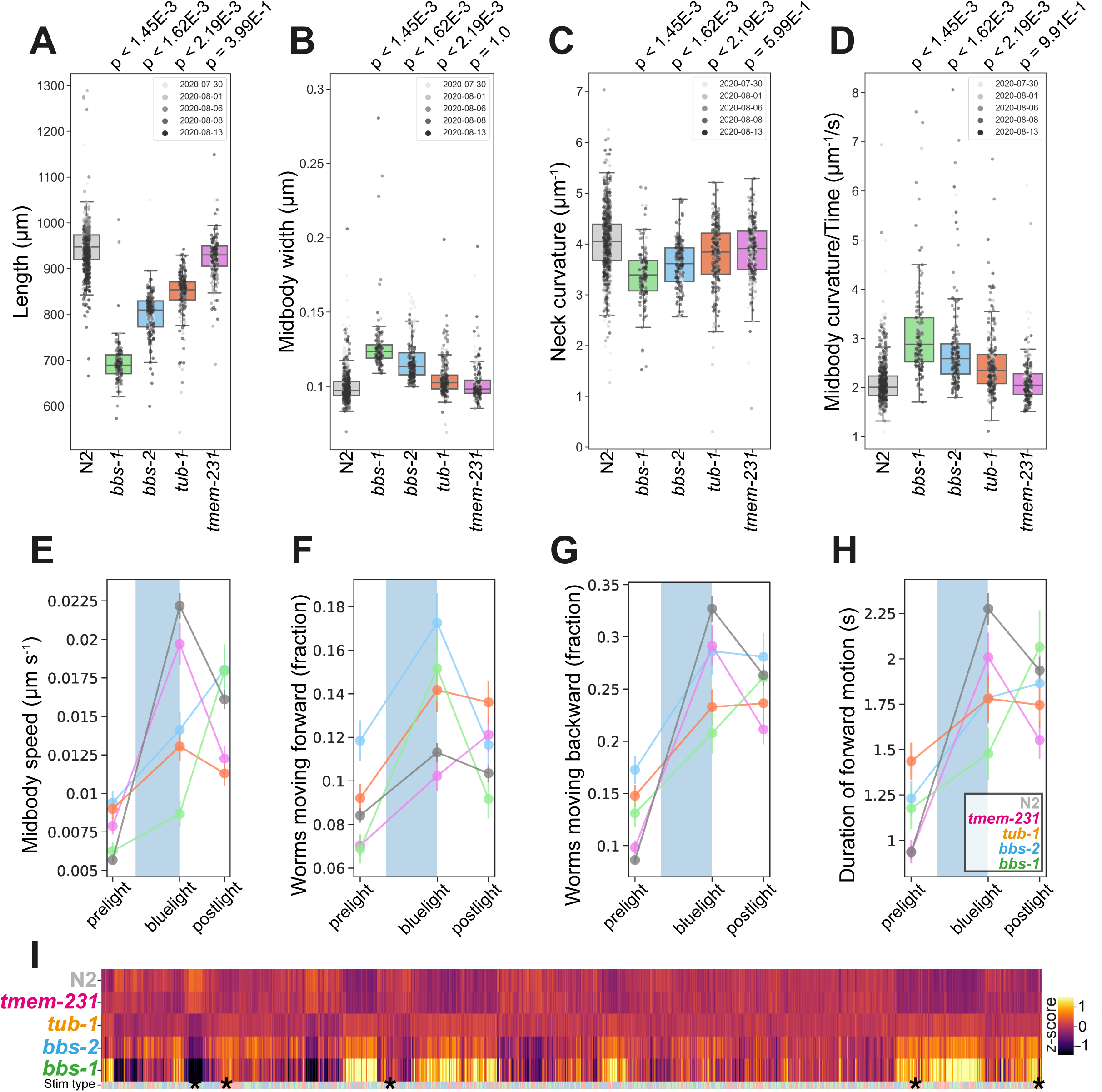
Ciliopathies disease model phenologs. **(A-D)** Key behavioural features altered in loss-of-function mutant strains associated with ciliopathies: *bbs-1(syb1588)*, *bbs-2(syb1547)*, *tub-1(syb1562)* and *tmem-231(syb1575)*, under baseline (pre-stimulus) imaging conditions. Individual points marked on the box plots are well averaged values (three worms per well) for each feature across the independent days of tracking. *P*-values are for comparisons to wild-type N2 worms using block permutation t-tests (*n* = 100,000 permutations, correcting for multiple comparisons using the Benjamini-Yekutieli method). **(E-H)** Changes in selected features in response to stimulation with a single 10-second blue light pulse (blue shaded region). Feature values were calculated using 10 second windows centred around 5 seconds before, 10 seconds after, and 20 seconds after the beginning of each blue light pulse. **(I)** Heatmap of the entire set of 8289 behavioural features extracted by Tierpsy for the disease model strains associated with ciliopathies and N2. The ‘stim type’ barcode denotes when during image acquisition the feature was extracted: pre-stimulation (pink), blue light stimulation (blue) and post-stimulation (green). Asterisks show the location of selected features present in A-E.

The vertebrate family of tubby-like proteins (TUB, TULP) are required for correct GPCR localisation and intra-flagellar transport in primary cilia (Mukhopadhyay & Jackson, 2011). In humans, mutations in this family are associated with rod-cone dystrophy, obesity, and retinitis pigmentosa (North et al., 1997). Furthermore, *Tub1-/-* mice reflect the associated human disease pleiotropy, with insulin resistance, obesity, and cochlea and visual degeneration (Sun et al., 2012). *C. elegans* have a single ortholog of tubby-like proteins, *tub-1*, and mutants have defects in chemotaxis and insulin signalling as well as increased lipid accumulation (Mak et al., 2006).

*tub-1(syb1562)* mutants are shorter, wider, and hyperactive compared to wild-type worms, similar to *bbs-1* and *bbs-2* mutants (Figures 3A, 3B and 3E). They also have a defect in their response to blue light (Figures 3F-H). Unlike BBS mutants, cilia integrity is maintained in *tub-1* loss-of-function mutants (Mak et al., 2006), which could explain the similar but less severe phenotype of *tub-1* mutants compared to *bbs-1* and *bbs-2* mutants.

Compartmentalisation of cilia-specific signalling components is regulated by a complex of transmembrane proteins including TMEM231 and mutations in this compartmentalisation complex are associated with neurodevelopmental limb defects and pathologies of the brain and kidneys found in Joubert and Meckel syndrome (Chih et al., 2012). Mouse *Tmem231-/-* mutations are embryonic lethal and the *C. elegans* ortholog *tmem-231* has been shown to have conserved molecular and cellular function (Roberson et al., 2015). However, unlike the other cilia-related mutants, *tmem-231* mutants do not have significant differences in body morphology or posture compared to N2 (Figure 3). We do note that *tmem-231(syb1575)* is slightly more active during baseline tracking (see strain-specific gene card).

### Channelopathies

The Na+ cation leak channel (NALCN) is expressed throughout the central nervous system, and in parts of the endocrine (pancreas, adrenal, thyroid gland), respiratory and cardiac systems (Cochet-Bissuel et al., 2014). NALCN is a voltage-independent, nonselective cation channel that regulates resting potential (Senatore et al., 2013), and plays a role in neuromodulation by neurotransmitters (Cochet-Bissuel et al., 2014). Mutations in NALCN are associated with neuromuscular disorders including severe hypotonia, infantile neuroaxonal dystrophy (INAD), congenital contractures, cognitive delay, autism, epilepsy, bipolar disorder, and cardiac/respiratory problems (Aoyagi et al., 2015; Cochet-Bissuel et al., 2014; Gal et al., 2016). NALCN-knock-out mice die 12 hours after birth (Lu et al., 2007), so development of appropriate non-mammalian animal models to understand the associated human diseases is essential.

*C. elegans* encodes two functionally redundant, but differentially expressed, NALCN homologs, NCA-2 and UNC-77, whose proper expression and axonal localization in cholinergic neurons is regulated by UNC-80 (Zhou et al., 2019). Unlike murine models, *nca-2*, *unc-77*, and *unc-80* loss-of-function mutants are viable. Similar to *Drosophila melanogaster* or mouse neonates lacking the cation leak channel (Cochet-Bissuel et al., 2014), loss-of-function mutations in *nca-2* and *unc-80* result in morphological changes in *C. elegans*: both mutants are significantly shorter than wild-type N2 (Figure 4A).

**Figure 4:**
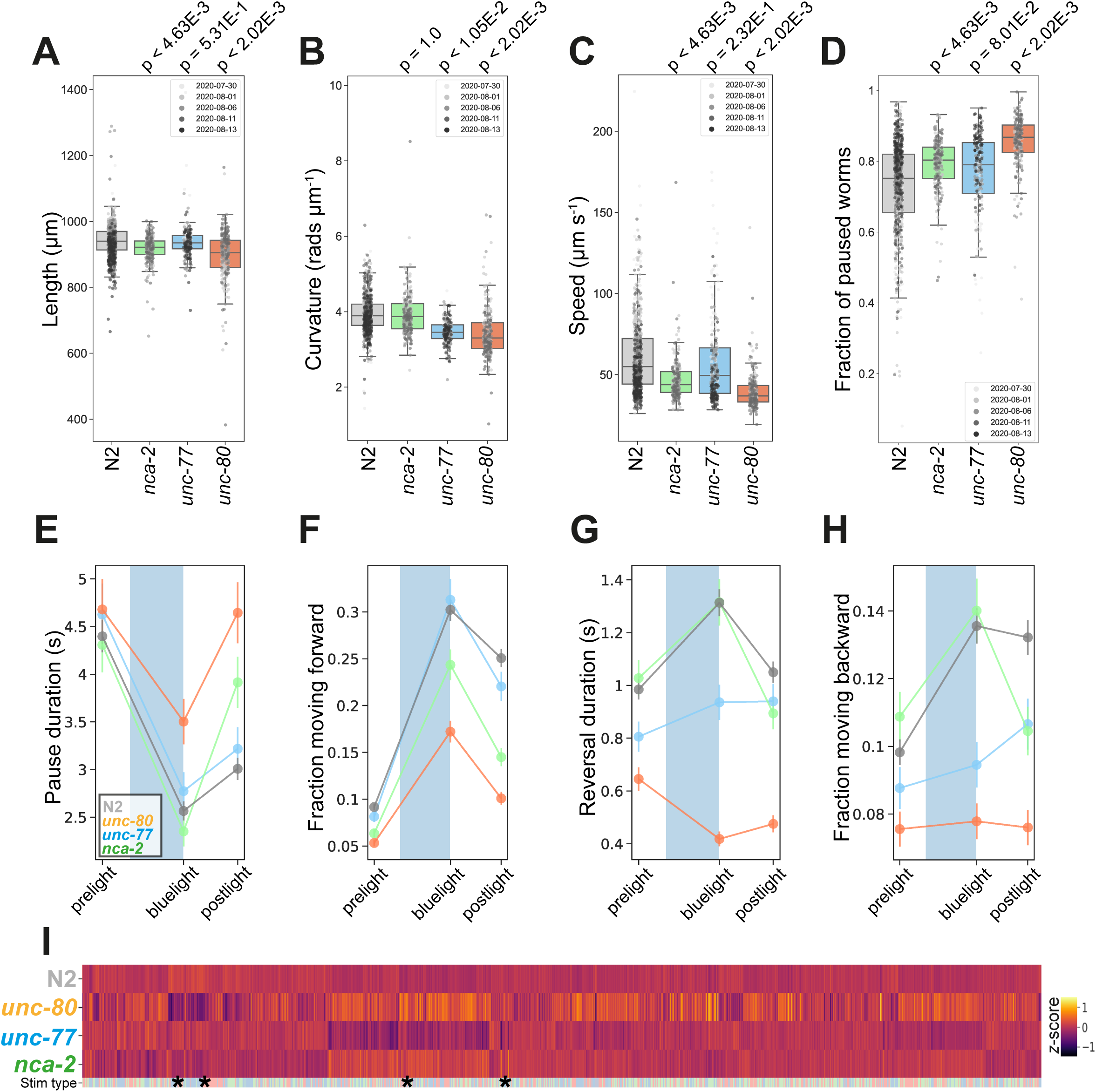
NALCN disease model phenologs. **(A-D)** Key behavioural and postural features altered in loss-of-function mutant strains associated with NALCN mutants: *nca-2(syb1612)*, *unc-77(syb1688)* and *unc-80(syb1531)*, under baseline (pre-stimulus) imaging conditions. Individual points marked on the box plots are well averaged values (three worms per well) for each feature across the independent days of tracking. *P*-values are for comparisons to wild-type N2 worms using block permutation t-tests (*n* = 100,000 permutations correcting for multiple comparisons using the Benjamini-Yekutieli method). **(E-H)** Changes in selected features in response to stimulation with a single 10-second blue light pulse (blue shaded region). Feature values were calculated using 10 second windows centred around 5 seconds before, 10 seconds after, and 20 seconds after the beginning of each blue light pulse. **(E)** A representative ‘fainting phenotype’ for *unc-80(syb1531)* and *nca-2(syb1612)*, characterised by an increase in pausing following the cessation of stimulation with blue light. **(I)** Heatmap of the entire set of 8289 behavioural features extracted by Tierpsy for the disease model strains associated with NALCN disease and N2. The ‘stim type’ barcode denotes when during image acquisition the feature was extracted: pre-stimulation (pink), blue light stimulation (blue) and post-stimulation (green). Asterisks show the location of selected features present in A-D.

Consistent with their differential expression (Jospin et al., 2007; Yeh et al., 2008), we observe phenotypic differences between *nca-2(syb1612)* and *unc-77(syb1688)* deletion mutants (Figure 4). Gain-of-function mutations in *unc-77* have previously been reported to cause deeper body bends (Topalidou et al., 2017) whereas *unc-77(syb1688)* deletion mutants have decreased curvature (Figure 4B). In contrast, *nca-2(syb1612)* has no significant change in curvature, but is slower than the wild-type strain (Figures 4B-C). We find that *unc-80(syb1531)* mutants, which affect the localisation of NALCN channel subunits, have the most severe phenotype exhibiting a decrease in both curvature and speed (Figures 4B-4C).

Mutations in both NALCN channel subunits in *C. elegans* cause a ‘fainter’ phenotype when stimulated mechanically or immersed in liquid (Pierce-Shimomura et al., 2008). Similarly, we found that a ten second blue light pulse resulted in increased post-exposure pausing and a decreased forward escape response in *nca-2(syb1612)* and *unc-80(syb1531)* single mutants (Figures 4E-H). *unc-77(syb1688)* mutants did not ‘faint’ after blue light exposure, but consistent with findings that the NALCN channelsome regulates reversal behaviour (Zhou et al., 2019) both *unc-77(syb1688)* and *unc-80(syb1531)* fail to initiate a backwards escape response. Pausing and ‘fainting’ after blue light exposure represents a novel screenable phenotype for NALCN-channelopathies.

### Drug repurposing screen in unc-80 mutants

Because *unc-80* mutants have a clear behavioural phenotype without a strong developmental phenotype, we reasoned that they were a useful test case for a drug repurposing screen using an acute 4-hour treatment which might rescue behavioural ‘symptoms’ in fully developed animals. When screening compound libraries, it is not practical to perform a large number of replicates which would be necessary to detect subtle phonotypes and overcome the reduced statistical power that comes from correcting for multiple comparisons in a large feature set. We therefore defined a reduced set of core features to capture the *unc-80* behavioural phenotype consisting of curvature, speed, and fraction of paused worms all during blue light stimulation (Figure 4B-D).

For the repurposing screen, we used a library of 743 FDA approved drugs. We prepared tracking plates where each well contained a drug at a final concentration of 100 μM with 1% DMSO. Three adult worms were added to each well, incubated for 4 hours, and then tracked. The screen was repeated across three independent tracking days with three independent wells per day (9 total well replicates). Hits were defined as compounds that significantly improved all three of the core *unc-80* features, meaning that treated worms were significantly different from *unc-80* DMSO controls and that the direction of the effect was towards wild-type controls. These hit compounds shift *unc-80* towards N2 in a three-dimensional phenotype space (Figure 5A). To empirically test our ability to detect hits when including more features in subsequent screens, we also repeated this analysis using pre-defined Tierpsy feature sets (Javer, Ripoll-Sánchez, et al., 2018) containing an increasing number of behavioural features. We found a clear reduction in statistical power when correcting for multiple comparisons across more behavioural features, with no hits being detected using >16 behavioural features (Supplementary Figure 1).

**Figure 5:**
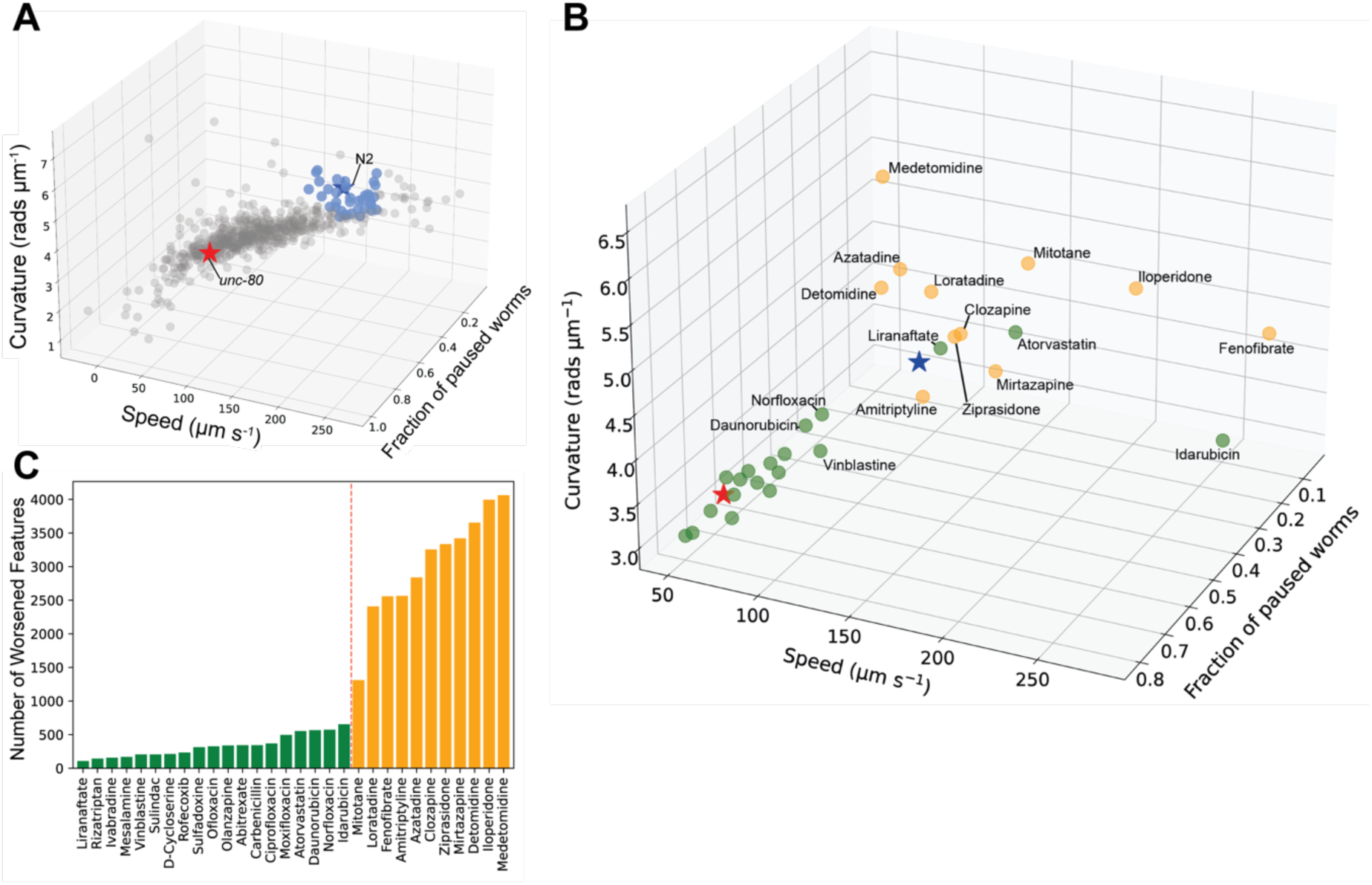
Drug repurposing screening. **(A)** Phenotypes of unc-80(syb1531) mutant (red star) and N2 (blue star) worms treated with 1% DMSO, and unc-80(syb1531) mutants treated with a library of 743 FDA approved drugs at a concentration of 100 μM for 4 hours (circles). Each point represents an average of 3 well replicates, across 3 independent days of tracking (n = 9 total). Blue points are the top 30 compounds that significantly improved all three of the core behavioural features, pushing the unc-80 mutant strain towards the control in phenospace. **(B)** Confirmation screen of the top 30 compounds identified in the initial library screen. Again unc-80(syb1531) and N2 DMSO treated controls are represented by red and blue stars, respectively, and each circular point represents unc-80(syb1531) treated with 100 μM compound for 4 hours. The 13 compounds coloured in yellow lead to the worsening of >1000 behavioural features (see below). Liranaftate and atorvastatin both lead to a rescue of the core mutant phenotype with a low number of side effects. **(C)** Total number of behavioural ‘side effects’ following treatment of unc-80(syb1531) with the 30 compounds in the confirmation screen. Side effects are defined as features that are not significant between unc-80 mutants and wild-type N2 worms treated with 1% DMSO but where there is a significant difference between unc-80 mutants treated with a drug compared to N2. Red dashed line separates drug treatments that lead to a worsening of >1000 behavioural features that correspond to the points coloured in yellow in the 3D scatterplot.

We then performed a confirmation screen on 30 of the hit compounds from the initial screen with a larger number of replicates (24 wells per day over three independent days). 13 of the 30 hits were reproduced in the sense of showing significant differences from *unc-80* DMSO controls in the direction of wild-type, however several showed a reduced effect size and were clustered around the *unc-80* DMSO controls (Figure 5B). Having a larger number of replicates, we also tested for ‘side effects’ in treated worms which we define as features where there was no significant difference between *unc-80* mutants and wild-type N2 animals but where there is a significant difference between *unc-80* mutants treated with a drug compared to wild-type N2 animals. About half of the confirmed hit compounds lead to more than 1000 side effects (Figure 5C) and are coloured yellow in the core *unc-80* phenotype space (Figure 5B), we also note that the same trend persists when looking at a reduced behavioural feature set of 256 predefined key Tierpsy features: the “Tierpsy 256” set (Javer, Ripoll-Sánchez, et al., 2018) (Supplementary Figure 2).

Of the confirmed hit compounds with fewer than 1000 side effects, liranaftate and atorvastatin (Lipitor) lead to good rescue of the core *unc-80* behavioural phenotypes. Interestingly, both compounds act upon the melavonate biosynthesis pathway. The main trunk of this pathway is conserved across species and converts acetyl-CoA to farnesyl diphosphate (Rauthan & Pilon, 2011). Atorvastatin is an inhibitor of HMG-CoA reductase that regulates lipid homeostasis (Goldstein et al., 2006) and catalyses the rate limiting step of the melavonate pathway. Liranaftate, an antifungal inhibitor of squalene epoxidase, shows the greatest behavioural improvement, with the fewest number of side effects. However, *C. elegans* lack the branch of the mevalonate pathway responsible for converting squalene to cholesterol, instead relying on dietary cholesterol. In this case the effect may be due to an off-target interaction on one of the other enzymes in the pathway.

Fenofibrate, a PPAR-alpha activator that increases lipolysis, is also a hit but has a large number of side effects. Given that it overshoots perfect rescue by shifting *unc-80* animals past wild-type controls in core phenospace, it could be that at a lower dose it would provide good rescue with fewer side effects. Since UNC-80 is involved in localisation of the NACLN+ channel complex, we hypothesise that changes in membrane composition might be indirectly affecting the localisation or function of channel complex.

## Discussion

We framed the problem of disease modelling as one of sampling and elucidating a model organism genotype-phenotype map that is connected to a human disease gene-phenotype map through genetic conservation. However, both the genomic and phenomic spaces are large and high-dimensional and so it was not clear that a systematic approach would identify useful phenologs for disease modelling. Using an initial set of genes that are diverse in function (but biased towards roles in neurons and muscles) and that lead to diverse symptoms when mutated in humans, we found that it is possible to systematically create and phenotype worm models of rare diseases using a uniform assay and protocol. Without adding additional perturbations or experimental conditions, we found that 23 out of the 25 disease models we made had detectable differences compared to wild-type.

Of the disease models with detectable phenotypes, approximately half had strong phenotypes that would support high throughput phenotypic screens for candidate treatments. To test the feasibility of performing drug repurposing screens using the same uniform assay, we chose a worm model of UNC80 deficiency and, focussing on a reduced core phenotype, identified an initial list of 30 FDA-approved drugs that rescued the mutant phenotype. A confirmation screen with more replicates identified several compounds that still rescued the core disease model phenotype and did not have a large number of side effects. The detection of drug side effects will be useful for prioritising hits for further follow-up by identifying nuisance compounds. For example, if a disease model is hyperactive and gross motility is the only measure used in a screen, compounds that are broadly toxic may be detected as hits but not be good drug candidates.

Our treatment approach of a short exposure time (4h) was chosen to identify compounds with a potential to rescue neural phenotypes once the developmental effects of a mutation have already occurred, mimicking clinical intervention of a pre-existing genetic disorder. However, the flexibility afforded by our standardised phenotyping methodology ensures it is possible to test longer, or shorter, periods of drug treatment. For example, L1s could be reared on drug-containing plates for their entire life before phenotyping. We note that this approach will be particularly well suited to identify compounds that rescue developmental delayed strains, such as of some of the disease model mutants used in this study (e.g., *cat-4*, *dys-1*, *figo-1*, *gpb-2* and *kcc-2* LoF mutants).

With a current discovery rate of >100 new genetic diseases each year, the number of diseases with little aetiological understanding is going to grow at a faster rate than biomedical and pharmaceutical research will find therapies (Boycott et al., 2017). Therefore, novel approaches that can systematically screen the effects of large numbers of genetic variants associated with different diseases are required in order to scale up drug discovery across all rare diseases. CRISPR genome editing in model organisms including *C. elegans* allows the creation of new disease models at the required rate and cost. For this proof-of-principal study, we focused on the generation of large deletion alleles to ensure putative LoF of the target gene. However, we note that a limitation of this approach may be the removal of regulatory genetic elements, e.g., non-coding RNAs or introns, as part of the deletion. For future screening efforts, particularly those against a single/smaller panel of genes, we suggest the generation and testing of additional LoF mutants through targeting functional domains or the introduction of early stop codons within a target allele. Better yet, the generation of specifically engineered alleles that match to a corresponding human disease condition will permit the generation of individual *C. elegans* “patient avatars” of diseases that will enable the development and testing of personalised therapeutics against patient-specific variants of genetic disorders. We have recently proven that our tracking approach is capable of separating individual *sod-1* mutations, associated with the onset of ALS in humans, in phenospace (Barlow et al., 2022). In summary, high-throughput behaviour recording with multidimensional phenotyping brings an *in vivo* drug repurposing screen within reach for every Mendelian disease caused by mutations in a conserved gene.

## Materials and Methods

### Mutant generation

CRISPR guide RNAs were designed to target large deletions (>1000bp) that start close to the start codon and excise several exons from the gene in order to give high confidence of loss of function. For exceptionally large genes (e.g. *dys-1* or *mpz-1*) the entire protein-coding region was excised. Mutants were designed and made by SunyBiotech in an N2 background.

### Worm preparation

All strains were cultured on Nematode Growth Medium at 20°C and fed with *E. coli* (OP50) following standard procedure (Stiernagle, 2006). Synchronised populations of young adult worms for imaging were cultured by bleaching unsynchronised gravid adults, and allowing L1 diapause progeny to develop for 2.5 days at 20°C (detailed protocol: https://dx.doi.org/10.17504/protocols.io.2bzgap6). Several strains were developmentally delayed and were allowed to grow for longer before imaging. *cat-4*(*syb1591*), *gpb-2*(*syb1577*), *kcc-2*(*syb2673*), and *unc-25*(*syb1651*) were allowed to develop for 3.5 and *dys-1*(*syb1688*), *figo-1*(*syb1562*), and *pink-1*(*syb1546*) were allowed to develop for 5.5 days prior to imaging. On the day of imaging, young adults were washed in M9 (detailed protocol: https://dx.doi.org/10.17504/protocols.io.bfqbjmsn), transferred to the imaging plates (3 worms per well) using a COPAS 500 Flow Pilot (detailed protocol: https://dx.doi.org/10.17504/protocols.io.bfc9jiz6), and returned to a 20°C incubator for 3.5 hours. Plates were then transferred onto the multi-camera tracker for another 30 minutes to habituate prior to imaging (detailed protocol: https://dx.doi.org/10.17504/protocols.io.bsicncaw).

For drug repurposing experiments, the MRCT FDA-approved compound library (Catalog No.L1300) was supplied pre-dissolved in DMSO by LifeArc (Stevenage, UK). The day prior to tracking, imaging plates were dosed with the compound library to achieve a final well concentration of 100 μM prior to seeding with bacteria (see below for details). Plates were left to dry (∼30 mins), before being stored in the dark at room temperature overnight. Following the methods described above, age synchronized young adult *unc-80*(*syb1531*) worms were dispensed into the imaging plate wells and incubated at 20°C for 4 hours before tracking. The behaviour of mutant worms dosed with the drugs was then compared to wild-type N2 and *unc-80*(*syb1531*) worms (also age-synchronized young adults) dispensed into the wells of the same tracking plates dosed with an identical volume (1% w/v) of DMSO only (detailed protocol: https://dx.doi.org/10.17504/protocols.io.5jyl8p5yrg2w/v1).

### Plate preparation

Low peptone (0.013%) nematode growth medium (detailed protocol: https://dx.doi.org/10.17504/protocols.io.2rcgd2w) was prepared as follows: 20 g agar (Difco), 0.13 g Bactopeptone, and 3 g NaCl were dissolved in 975 mL of milliQ water. After autoclaving, 1 mL of 10 mg/mL cholesterol was added along with 1 mL CaCl2 (1 M), 1 mL MgSO4 (1 M) and 25 mL KPO4 buffer (1M, pH 6.0). Molten agar was cooled to 50-60°C and 200 μL was dispensed into each well of 96-square well plates (Whatman UNIPLATE: WHAT-77011651) using an Integra VIAFILL (detailed protocol: https://dx.doi.org/10.17504/protocols.io.bmxbk7in). Poured plates were stored agar-side up at 4°C until required.

One day prior to imaging, plates were placed without lids in a drying cabinet to lose 3-5% of weight by volume. Wells were then seeded with 5 μL OP50 (OD600 1.0) using an Integra VIAFILL dispenser, and plates were stored with lids (WHAT-77041001) on at room temperature overnight.

### Image acquisition

All videos were acquired and processed following methods previously described (Barlow et al., 2022). In brief, videos were acquired at 25 frames per second in a room with a nominal temperature of 20°C using a shutter time of 25 ms and a resolution of 12.4 µm/px. Three videos were taken sequentially: a 5-minute pre-stimulus video, a 6-minute blue light recording with three 10-second blue light pulses starting at 60, 160 and 260 seconds, and a 5-minute post-stimulus recording. The script for controlling recording timings and photostimulation was made using LoopBio’s API for their Motif software (https://github.com/loopbio/python-motifapi).

### Image processing and feature extraction

Videos were segmented and tracked using Tierpsy Tracker (Javer, Currie, et al., 2018). After segmentation of worm skeletons, our previously described convolutional neural network classifier was used to exclude non-worm objects from being classified during feature extraction (Barlow et al., 2022). Skeletons that did not meet the following criteria were removed from the analysis: 700 – 1300 µM length, 20 – 200 µM width. As an additional quality control measure, we used Tierpsy Tracker’s viewer to mark wells with visible contamination, agar damage, compound precipitation, or excess liquid as “bad”, and exclude these wells from downstream analysis.

Following tracking, we extracted a previously-defined set of 3076 behavioural features for each well in each of the three videos (pre-stimulus, blue light, and post-stimulus) (Javer, Ripoll-Sánchez, et al., 2018). Feature values are averaged over tracks to produce a single feature vector for each well.

### Statistical analysis

Statistically significant differences in the behavioural feature sets extracted by Tierpsy were calculated using block permutation t-tests in a pairwise manner between the mutant and/or treatment of interest and wild-type N2. Python (version 3.8.5) and the ‘univariate_tests’ function of Tierpsytools ‘statistical_tests’ module (https://github.com/Tierpsy/tierpsy-tools-python/blob/master/tierpsytools/analysis/statistical_tests.py) were used to perform these analysis. The *p*-value of each feature is calculated using *n* = 10000 permutations that randomly shuffle data labels within, but not between, the independent days of image acquisition for each strain/treatment to control for day-to-day variation in the technical replicates of each experiment. The resulting *p*-values were then corrected for multiple comparisons using the Benjamimi-Yekutieli procedure to control the false discovery rate at 5% (Benjamini & Yekutieli, 2001) and *p* > 0.05 is considered not statistically significant.

Heatmaps and principal component analysis show the *z*-normalised values of the extracted behavioural feature sets compared to the N2 reference and were calculated using in-built Seaborn (version 0.11.2) packages (Waskom, 2021). All the scripts used for statistical analysis and the generation of figures are available at: (https://github.com/Tom-OBrien/Systematic-creation-and-phenotyping-of-Mendelian-disease-models-in-C.elegans).

## Data Availability

The datasets (Tierpsy features, tracking data, and metadata) and code produced in this study are available on Zenodo, DOI: 10.5281/zenodo.12547170. Any remaining information can be obtained from the corresponding author upon reasonable request.

## Supporting information

Strain Specific Gene Cards

## Acknowledgements

This project has received funding from the European Research Council (ERC) under the European Union’s Horizon 2020 research and innovation programme (Grant agreement No. 714853) and was supported by the Medical Research Council through grant MC-A658-5TY30.

