## Supplementary material for "Systematic creation and phenotyping of Mendelian disease models in *C. elegans*: towards large-scale drug repurposing": Strain Specific Gene Cards

### *add-1(syb1598)*

Strain name: PHX1598
Wild-type gene length: 1951bp
Mutation: 1539bp deletion (second half of exon 1 to exon 5)

#### *C. elegans* Description

<u>Function</u>: Encodes cytoskeletal alpha-adducin; associated with cell membrane-mediated localised changes in cytoskeleton in response to activated signal transduction cascades

<u>Expression</u>: Coelomocytes; neurons; intestinal and rectal epithelia

<u>Previously reported phenotypes</u>: Defective short/long term memory and defective GLR-1 ionotropic glutamate receptor dynamics[1]

#### Human Orthologs:

*ADD1; ADD3*

<u>Associated disease(s):</u>
Artery disease; cerebral palsy, spastic quadriplegic, 3, autosomal recessive (AR) 617008; Hypertension, essential, salt-sensitive, mutational (Mu) 145500

#### Results

494/8289 significant features vs N2 ($p < 0.05$, block permutation t-test, 10000 permutations)

Key phenotype(s): increased curvature and angular velocity of the head

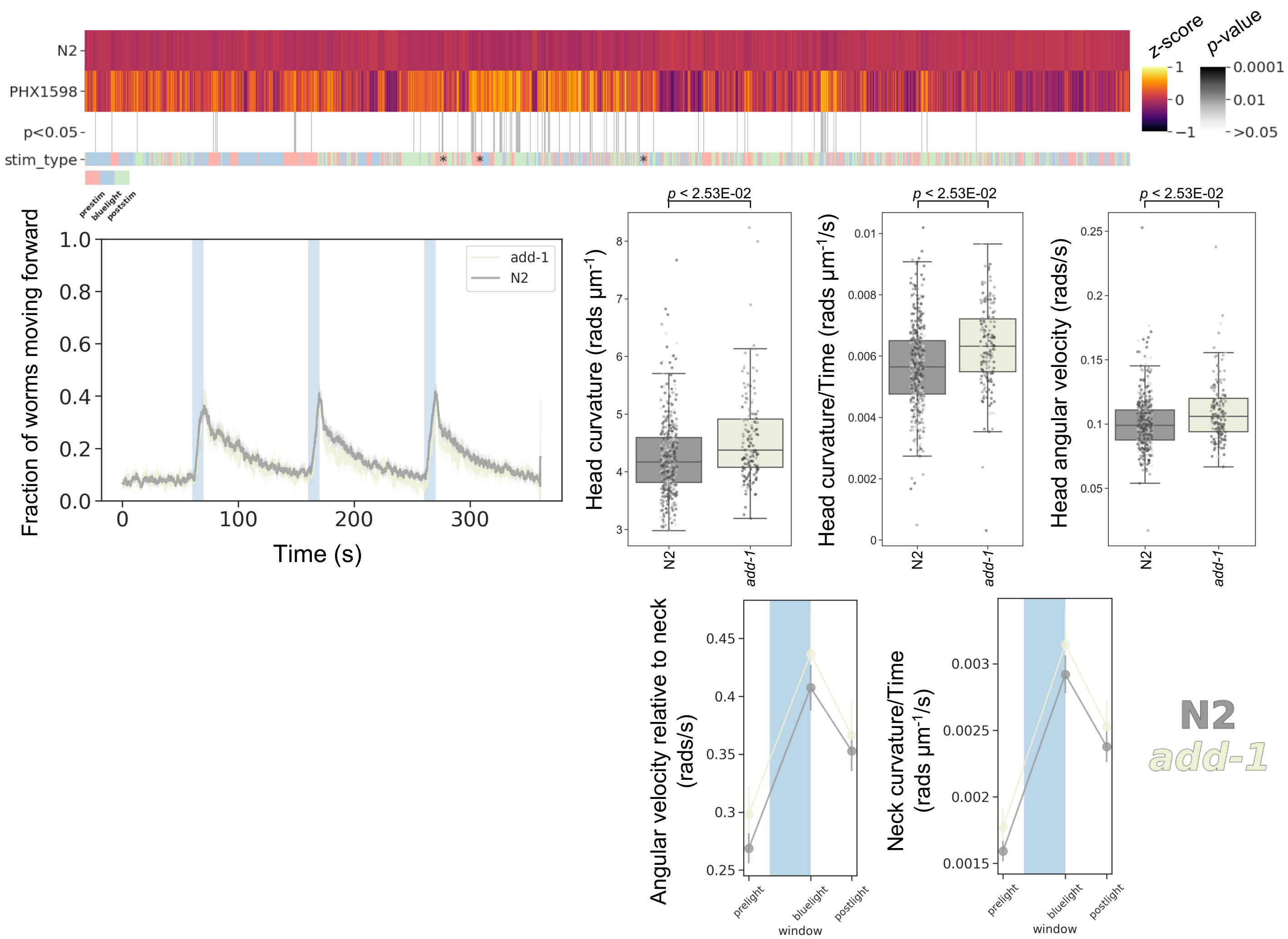

### ***avr-14(syb1576)***

Strain name: PHX1576
Wild-type gene length: 1111bp
Mutation: 628bp deletion (exon 1 and start of exon 2)

#### *C. elegans* Description

Function: Localises to membrane and encodes glutamate-gated chloride channel activity (ivermectin sensitive)

Expression: Neurons and somatic nervous system

Previously reported phenotypes: Defective salt-chemotaxis behaviour and decreased locomotion on food[2-3]

#### Human Orthologs:

*GLRA1, GLRB, GLRA3, GLRA4*

Associated disease(s):
Hyperekplexia, hereditary 1, autosomal dominant (AD) or recessive (AR), 149400; Hyperekplexia 2, 614619 (AR)

#### Results

2717/8289 significant features vs N2 ($p < 0.05$, block permutation t-test, 10000 permutations)

Key phenotype(s): small; increased curvature of the head and midbody; decreased head foraging (radial velocity); attenuated blue light response (e.g., decreased increase in speed upon stimulation with blue light)

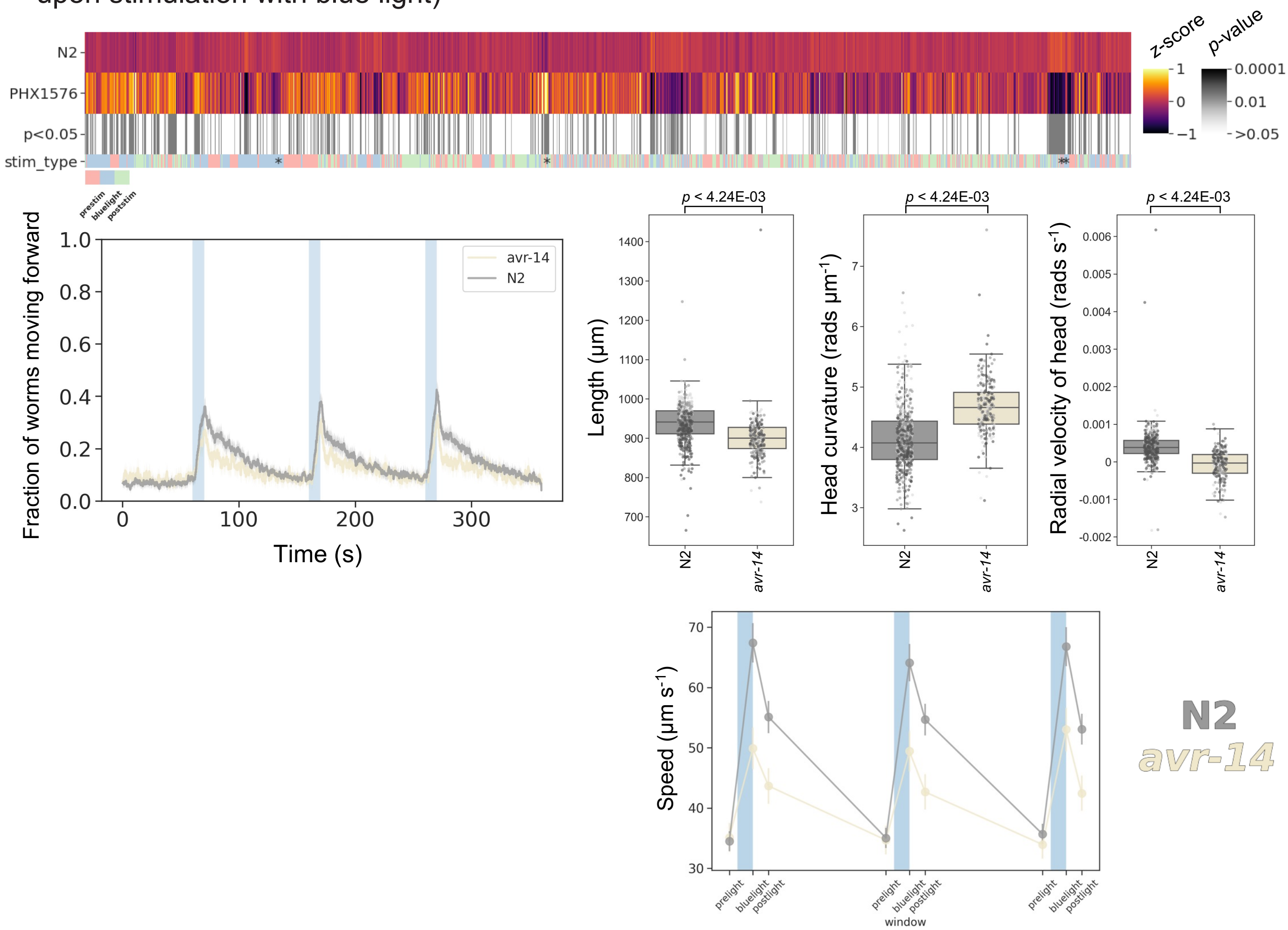

### ***bbs-1(syb1588)***

Strain name: PHX1588
Wild-type gene length: 1984bp
Mutation: 1412bp deletion (exon 1 to start of exon 2)

#### *C. elegans* Description

Function: Involved in intraciliary retrograde transport, non-motile cilium assembly and protein localization to cilium; regulates fat storage, as such, *bbs-1* affects body size, feeding, chemosensation and is required for proper regulation of insulin secretion

Expression: Ciliated neurons and sensory neurons

Previously reported phenotypes: Increased secretion of insulin, biogenic amines and neuropeptides; abnormal dye filling; ivermectin resistance[4-5]

#### Human Orthologs:

*BBS1*

Associated disease(s):
Bardet-Biedl syndrome 1, autosomal recessive (AR) or digenic recessive (DR), 209900

#### Results

6124/8289 significant features vs N2 ($p < 0.05$, block permutation t-test, 10000 permutations)

Key phenotype(s): shorter and fatter; decreased curvature; deeper body bends (time derivative of curvature); hyperactive prior to simulation; delayed, but sustained, forward escape response and an increased, but short-lived, backward escape response to blue light

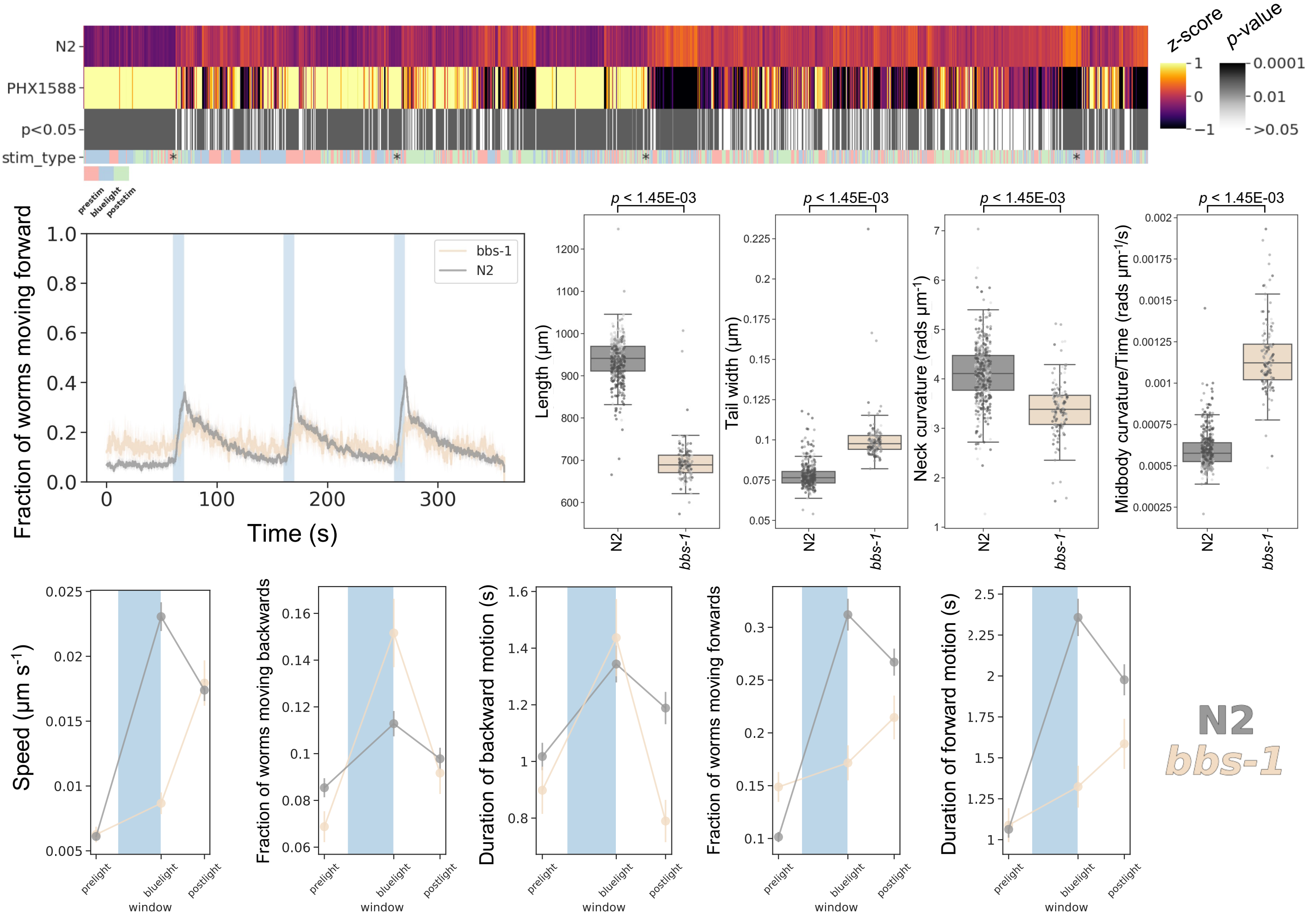

### ***bbs-2(syb1547)***

Strain name: PHX1547
Wild-type gene length: 2330bp
Mutation: 1513bp deletion (second half exon 1 to half of exon 4)

#### *C. elegans* Description

<u>Function</u>: Involved in non-motile cillum assembly

<u>Expression</u>: Ciliated neurons and sensory neurons

<u>Previously reported phenotypes</u>: Body size, developmental delay and roaming defects[4]

#### Human Orthologs:

*BBS2*

<u>Associated disease(s):</u>
Bardet-Biedl syndrome 2, 615981 (AR); Retinitis pigmentosa 74, 616562 (AR)

#### Results

5693/8289 significant features vs N2 ($p < 0.05$, block permutation t-test, 10000 permutations)

Key phenotype(s): shorter and fatter; decreased curvature; deeper body bends (time derivative of curvature); hyperactive prior to stimulation; increased (but delayed) increase in speed, short lived backward escape response, and delayed (but sustained) forward escape response to blue light stimulation

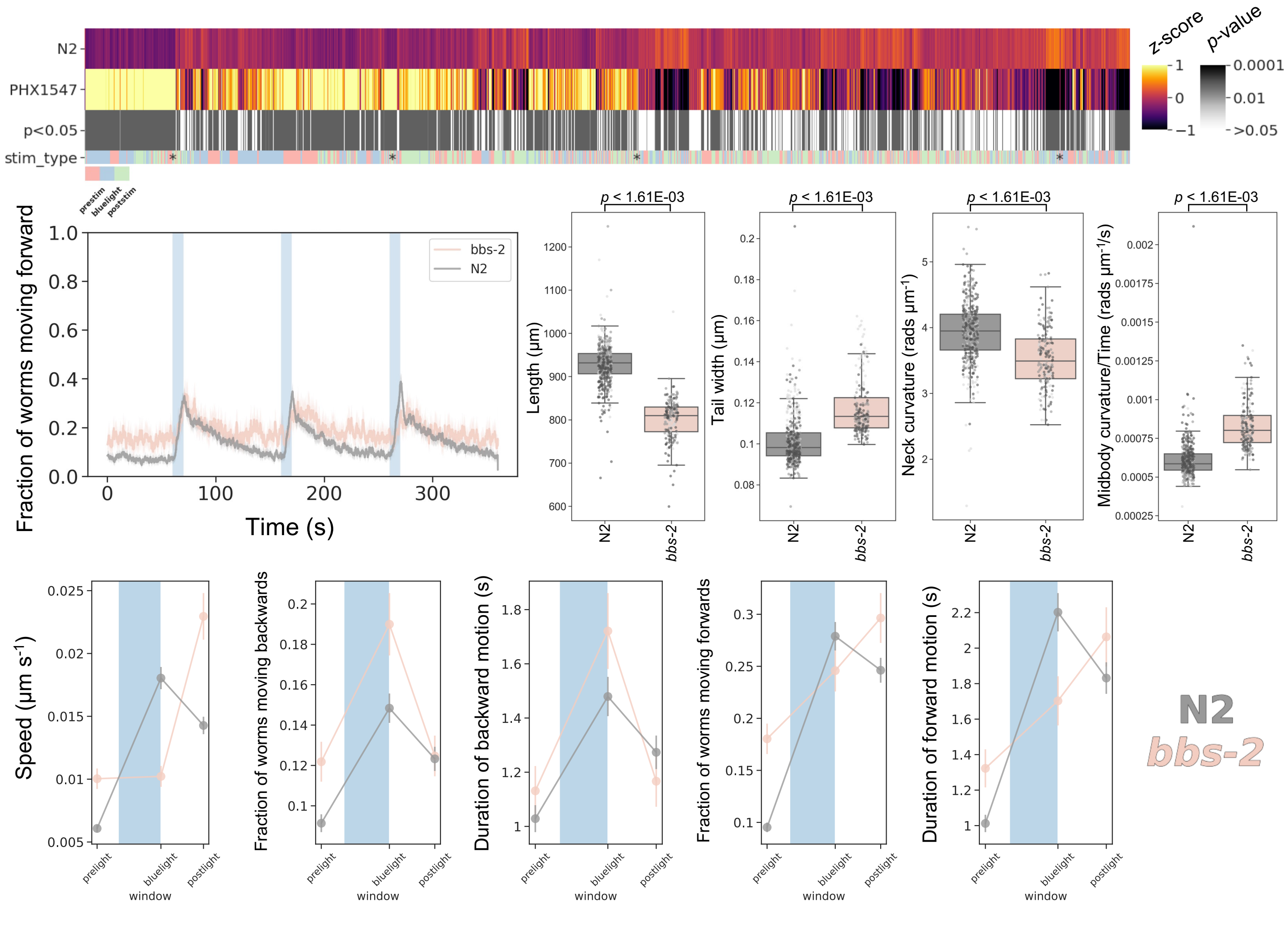

### ***cat-2(syb1542)***

Strain name: PHX1542
Wild-type gene length: 1683bp
Mutation: 1520bp deletion (half of exon 1 to half of exon 3)

#### *C. elegans* Description

Function: Tyrosine hydroxylase activity (catecholamine synthesis)

Expression: Dopaminergic neurons and male-specific anatomy (spiricules and ray neurons)

Previously reported phenotypes: Decreased dopamine levels; defective food sensing; increased peak acceleration; increased variation in speed; increased habituation in response to nonlocalized mechanical stimulation[6-7]

#### Human Orthologs:

*TH*

Associated disease(s):
*L*-Dopa-responsive dystonia/Segawa syndrome, 605407 (AR); Obesity; Parkinsonism

#### Results

4479/8289 significant features vs N2 ($p < 0.05$, block permutation t-test, 10000 permutations)

Key phenotype(s): increased speed/curvature of head tip and decreased head foraging (radial velocity of head) while moving forward; hypersensitive to blue light stimulation (e.g., greater increase in speed and angular velocity); short backward escape response to blue light

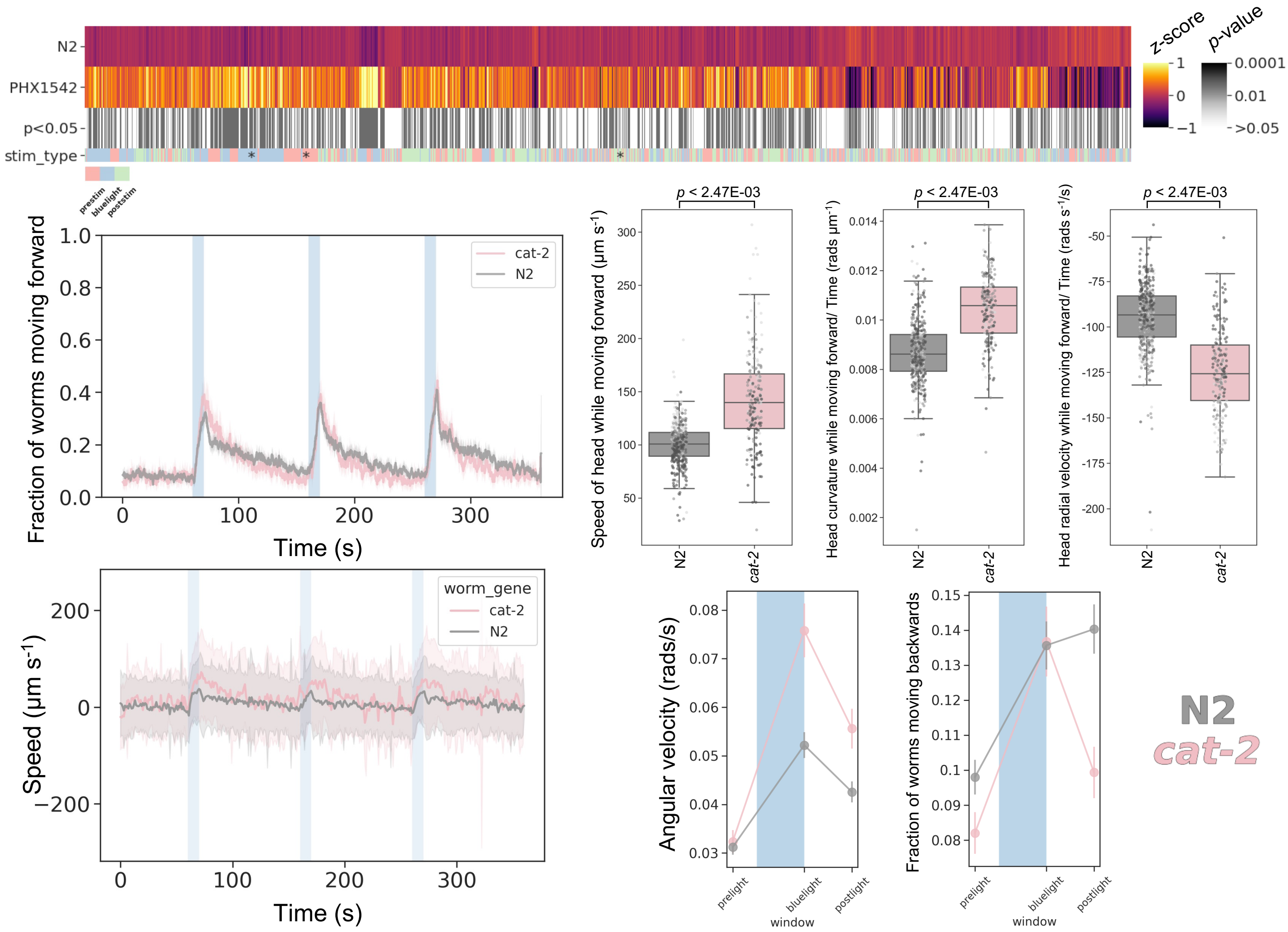

### ***cat-4(syb1591)***

Strain name: PHX1591
Wild-type gene length: 913bp
Mutation: 891bp deletion (exons 2-3 and half of exon 4)

#### *C. elegans* Description

Function: Putitative GTP cyclohydrolase I activity

Expression: Muscle cells; pharynx motor neurons; intestinal cells; dopaminergic, serotinergic, and mechanosensory neurons

Previously reported phenotypes: No previous phenotype reported

#### Human Orthologs:

*GHC1*

Associated diseases (from OMIM):
*L*-Dopa-responsive dystonia, with or without hyperphenylalaninemia, 128230 (AD, AR); Hyperphenylalaninemia, BH4 deficient, B, 233910 (AR); Bipolar disorder

#### Results

4182/8289 significant features vs N2 ($p < 0.05$, block permutation t-test, 10000 permutations)

Key phenotype(s): Developmental delay (2 days); small brood size; smaller; increased curvature; decreased speed; less active; strong photophobic escape response; short duration of backwards escape response

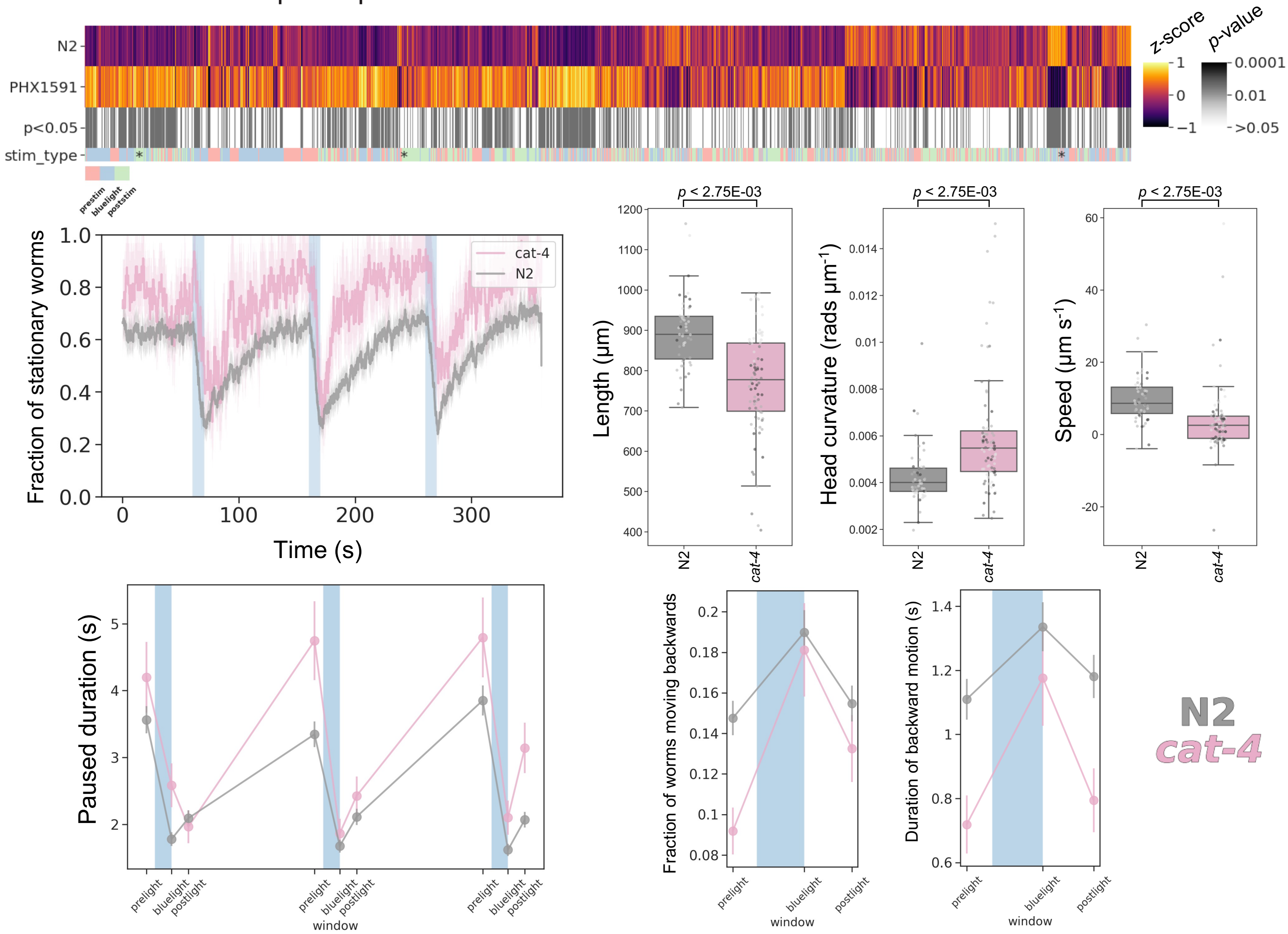

### *dys-1(syb1688)*

Strain name: PHX1688
Wild-type gene length: 30990bp
Mutation: 30716bp deletion (all of the F32B4.10 transcript)

#### *C. elegans* Description

<u>Function</u>: Localizes to dystrobrevin complex

<u>Expression</u>: Body wall musculature; head muscle; pharyngeal muscle; vulval muscle

<u>Previously reported phenotypes</u>: Subtle hyperactive locomotion; exaggerated head bending; slight hypercontractions; abnormal burrowing[8-9]

#### Human Orthologs:

*DMD*

<u>Associated disease(s):</u>
Duchenne muscular dystrophy, X-linked recessive (XLR) 310200; Becker muscular dystrophy, 300376 (XLR); Cardiomyopathy dilated, 302045 (XLR)

#### Results

1038/8289 significant features vs N2 ($p < 0.05$, block permutation t-test, 10000 permutations)

Key phenotype(s): Developmental delay (4 days); increased curvature, angular velocity and speed of head; increased sensitivity to blue light stimuli (e.g., greater increase in speed/forward motion upon stimulation); strongly attenuated backward escape response to blue light stimulation

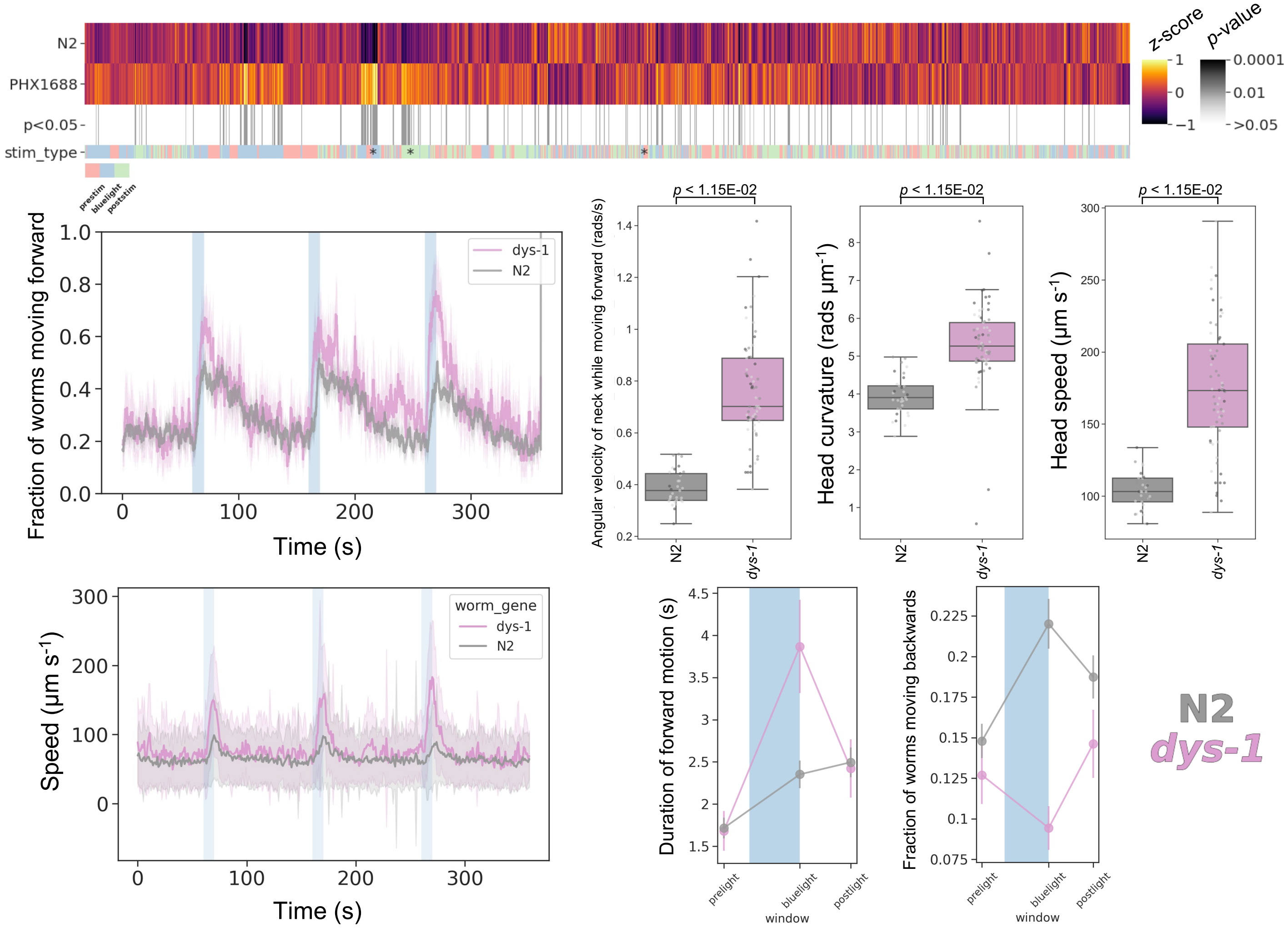

### ***figo-1(syb1562)***

Strain name: PHX1562
Wild-type gene length: 1633bp
Mutation: 1038bp deletion (first 3 exons)

#### *C. elegans* Description

<u>Function</u>: Putative phosphoric ester hydrolase activity

<u>Expression</u>: GABAergic neurons; body wall musculature; head neurons; intestine and tail neurons
<u>Previously reported phenotypes</u>: No previous phenotype reported

#### Human Orthologs:

*FIG4*

<u>Associated disease(s):</u>
Amyotrophic lateral sclerosis 11, 612577 (AD); Charcot-Marie-Tooth disease, type 4J, 611228 (AR); Polymicrogyria, bilateral temporooccipital, 612691 (AR); Yunis-Varon syndrome, 216340 (AR)

#### Results

2799/8289 significant features vs N2 ($p < 0.05$, block permutation t-test, 10000 permutations)

Key phenotype(s): Developmental delay (2 days); increased angular velocity/curvature of the head, neck and tail; deeper body bends (time derivative of curvature); sustained forward, but attenuated backward, escape response to blue light stimulation; greater increase in speed upon repeated blue light stimulation

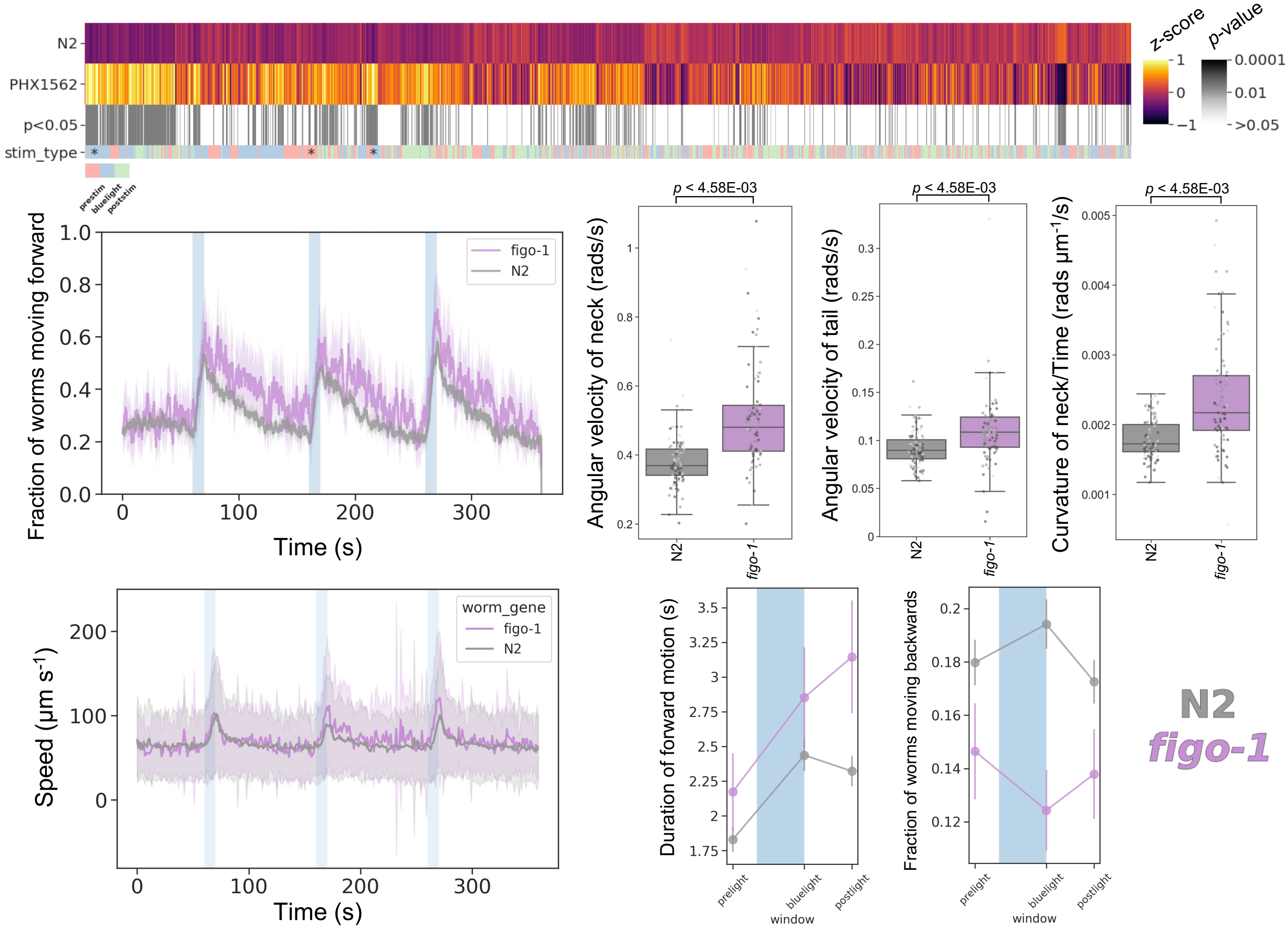

### *glc-2(syb1557)*

Strain name: PHX1557
Wild-type gene length: 1512bp
Mutation: 1269bp deletion (first 4 exons)

#### *C. elegans* Description

<u>Function</u>: Glutamate-gated Chloride channel (GLC-2), capable of forming homomeric glutamate-activated channels and heteromeric glutamate-activated channels with GLC-1; avermectins or anthelmintics inhibit pharyngeal pumping

<u>Expression</u>: Pharyngeal pm4 muscle

<u>Previously reported phenotypes</u>: No previous phenotype reported

#### Human Orthologs:

*GLRA1; GLRB*

<u>Associated disease(s):</u>
*Hyperekplexia, hereditary 1, 149400 (AD or AR); Hyperekplexia 2, 614619 (AR)*

#### Results

2413/8289 significant features vs N2 ($p$ < 0.05, block permutation t-test, 10000 permutations)

Key phenotype(s): Smaller; increased curvature and speed of the head; hyperactive; increased fraction and duration of forward motion; attenuated backward escape response; diminished blue light response (e.g., change in angular velocity upon stimulation is decreased)

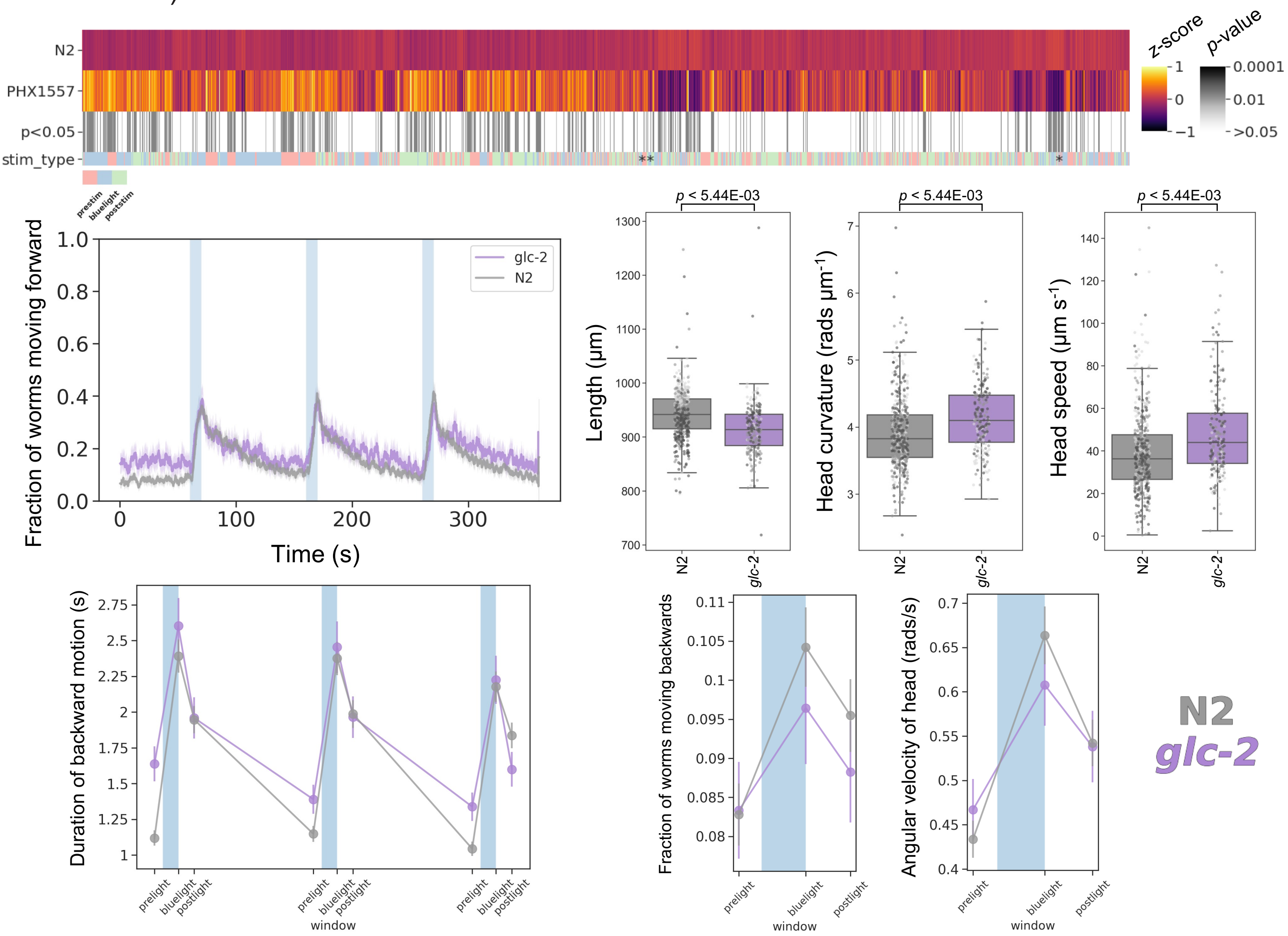

### *glr-1(syb1556)*

Strain name: PHX1556
Wild-type gene length: 1488bp
Mutation: 979bp deletion (exon 1 to beginning of exon 4)

#### *C. elegans* Description

Function: Localizes to several cellular components, including: ionotropic glutamate receptor complex; neuron projection membrane and perinuclear endoplasmic reticulum

Expression: Ganglia; neurons and ventral nerve cord

Previously reported phenotypes: Gain of function mutants are resistant to dopamine-induced food paralysis and have higher reversal rate; loss of function mutants are defective in light nose touch response; resistance to fluoxetine-induced paralysis[10-12]

#### Human Orthologs:

*GRIA3, GRIK2*

Associated disease(s):
Intellectual disability, 94, 300699 (XLR); Intellectual disability, 6, 611092 (AR); Huntington's

#### Results

3820/8289 significant features vs N2 ($p$ < 0.05, block permutation t-test, 10000 permutations)

Key phenotype(s): Longer; increased foraging; increased curvature/angular velocity of all body segments; increased speed; hyperactive; attenuated backward escape response to blue light

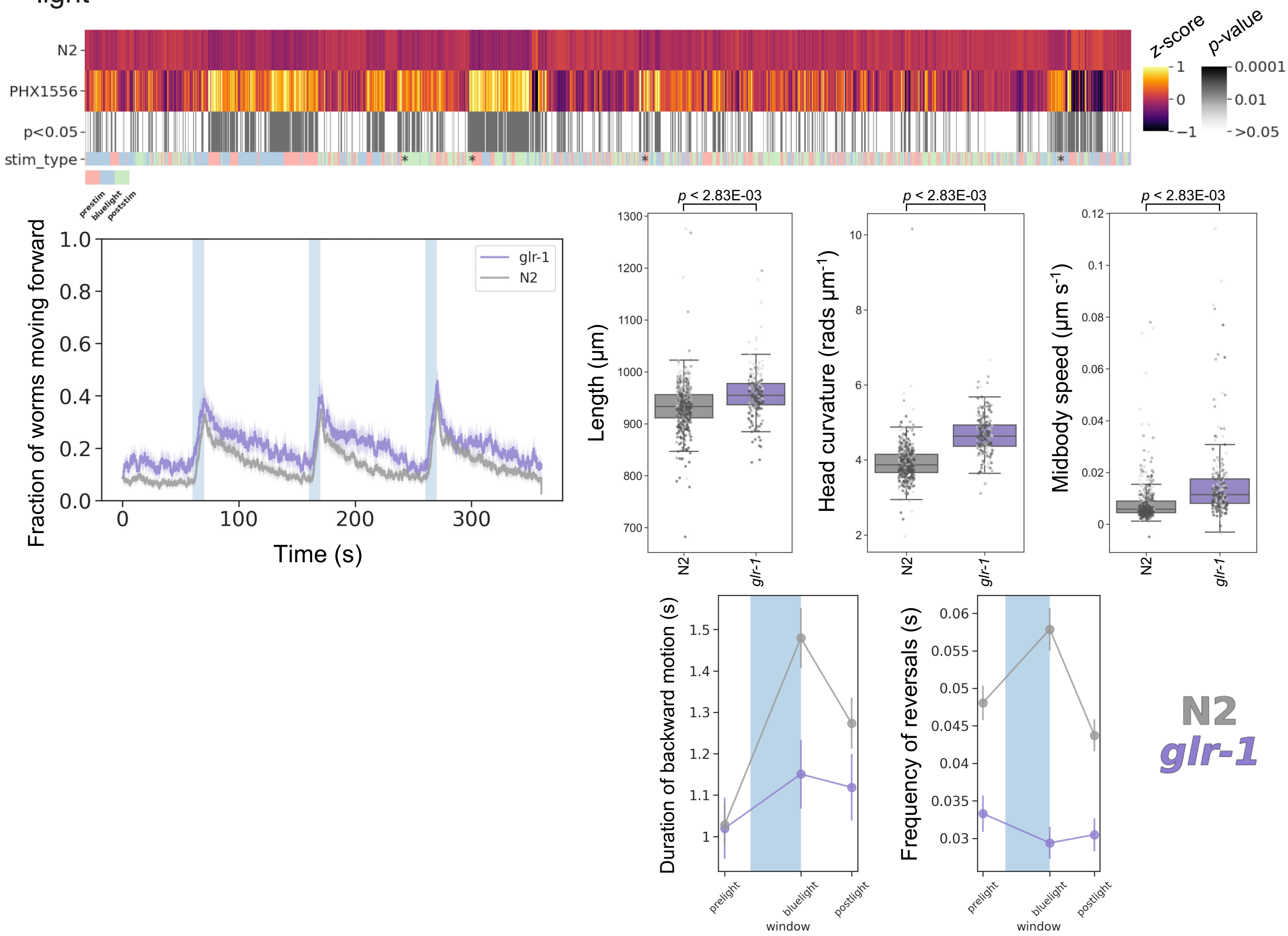

### *glr-4(syb1518)*

Strain name: PHX1518
Wild-type gene length: 2124bp
Mutation: 1356bp deletion (end of exon 1 to exon 8)

#### *C. elegans* Description

Function: NMDA-receptor subunit (encodes a putative non-NMDA ionotropic glutamate receptor subunit)

Expression: DA9; SABVL; SABVR and VB neurons

Previously reported phenotypes: Aldicarb resistance[13]

#### Human Orthologs:

*GRIK2*

Associated disease(s):
Intellectual disability, 6, 611092 (AR)

#### Results

0/8289 significant features vs N2 ($p < 0.05$, block permutation t-test, 10000 permutations)

Key phenotype(s): No significant phenotype detected

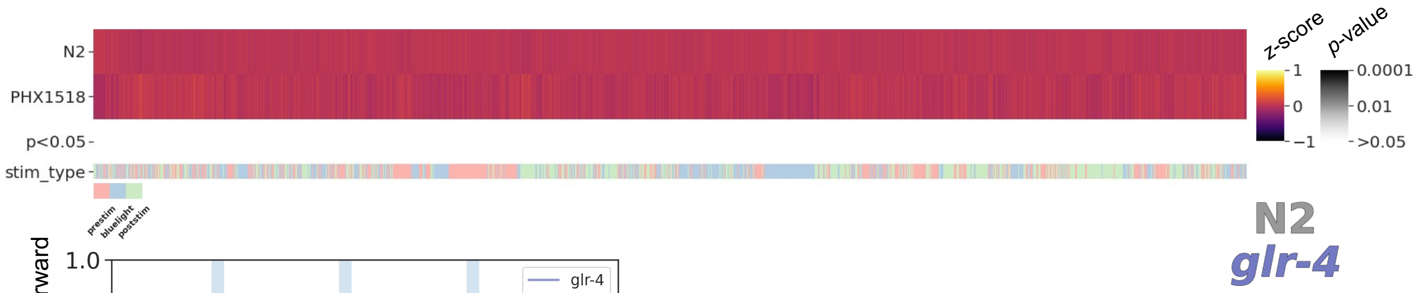


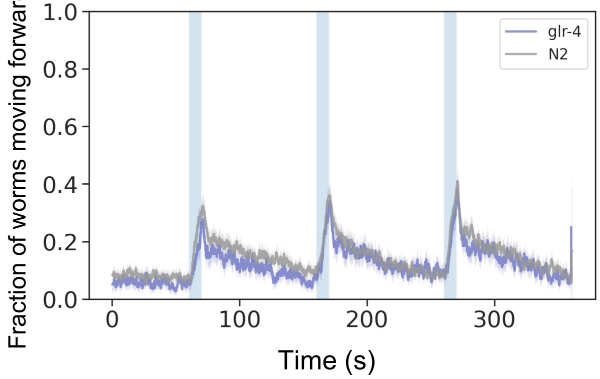

### *gpb-2(syb1577)*

Strain name: PHX1577
Wild-type gene length: 3009bp
Mutation: 2408bp deletion (all exons)

#### *C. elegans* Description

<u>Function</u>: G protein-coupled acetylcholine receptor signalling pathway; egg-laying

<u>Expression</u>: Body wall musculature; nerve ring; pharyngeal muscle cell and vulval muscle

<u>Previously reported phenotypes</u>: Slippery corpus; slightly longer; increased sinusoidal and deeper body bends; defects in egg laying and pharyngeal pumping; arecoline sensitive[14-16]

#### Human Orthologs:

*GNB5*

<u>Associated disease(s):</u>
Intellectual developmental disorder with cardiac arrhythmia, 617173 (AR); Language delay and ADHD/cognitive impairment with or without cardiac arrhythmia, 617182 (AR)

#### Results

2954/8289 significant features vs N2 ($p < 0.05$, block permutation t-test, 10000 permutations)

Key phenotype(s): Developmental delay (2 days); increased curvature/angular velocity of hips and tail; increased sinusoidal movements (time derivative of curvature); hyperactive; stronger, more sustained, backward escape response; hypersensitive to blue light

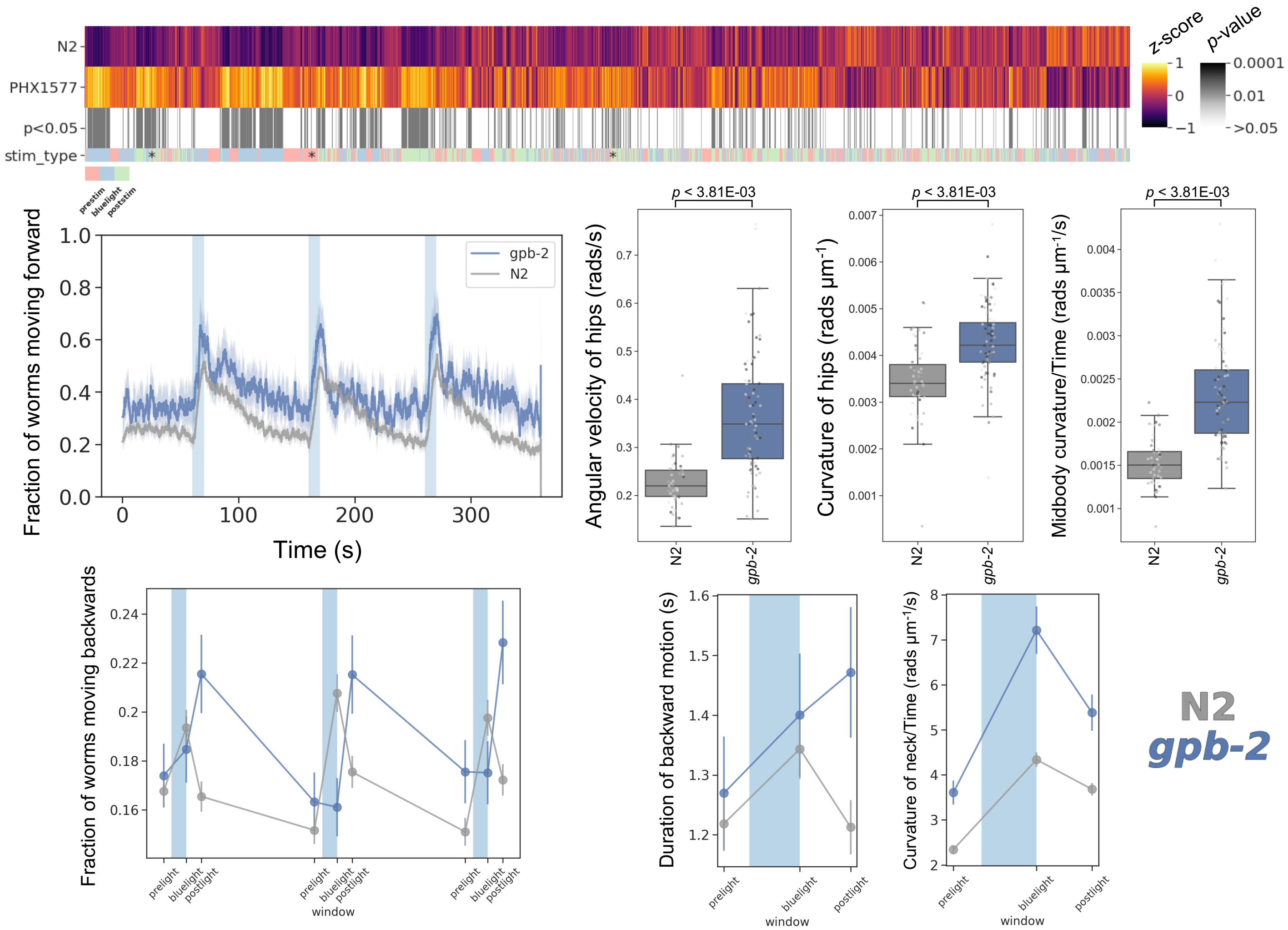

### *kcc-2(syb2673)*

Strain name: PHX2673
Wild-type gene length: 4387bp
Mutation: 3919bp deletion (first 3 exons and most piwi-interacting RNAs)

#### *C. elegans* Description

Function: Potassium-chloride c-transporter (partially redundant with abts-1); involved in chloride/potassium ion transport and regulation of GABAergic synaptic transmission

Expression: Body wall musculature; egg-laying apparatus; hermaphrodite distal tip cell; intestine and neurons

Previously reported phenotypes: Decreased egg laying; shorter; serotonin resistant; displays an excitatory, not inhibitory, response to the GABAa agonist muscimol[17-18]

#### Human Orthologs:

*SLC12A5, SLC12A6*

Associated disease(s):

Epilepsy, idiopathic generalized, 14, 616685 (AD); Epileptic encephalopathy, early infantile, 34, 616645 (AR); Agenesis of the corpus callosum with peripheral neuropathy, 218000 (AR)

#### Results

3246/8289 significant features vs N2 ($p$ < 0.05, block permutation t-test, 10000 permutations)

Key phenotype(s): Developmental delay (2 days); increased curvature and speed of head; decreased sinusoidal undulations (time derivative of midbody curvature); decreased angular velocity; hypoactive; sustained forward escape response; attenuated blue light response

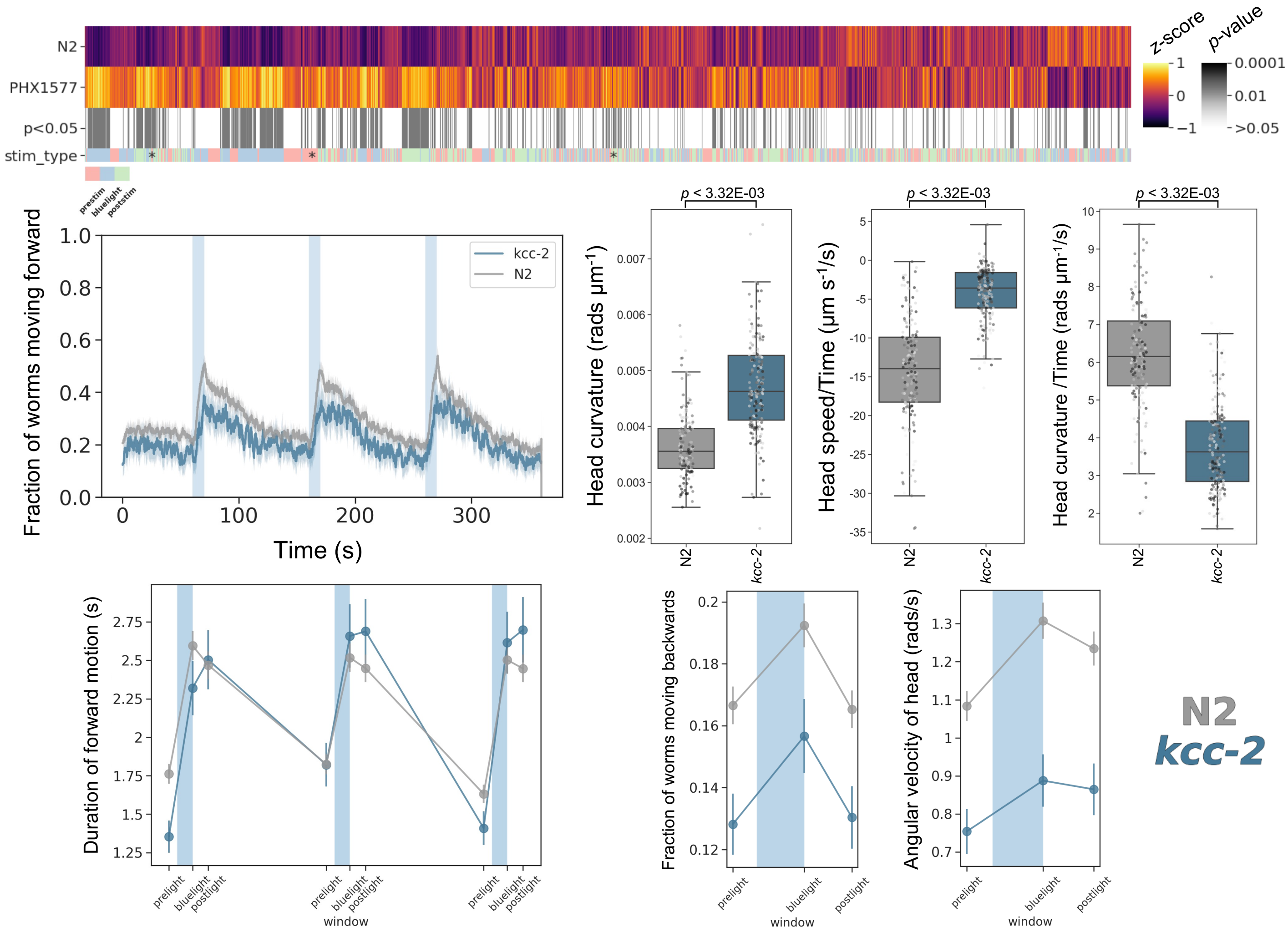

### ***mpz-1(syb1522)***

Strain name: PHX1522
Wild-type gene length: 36850bp
Mutation: 35787bp deletion (entire gene)

#### *C. elegans* Description

Function: Encodes multi-PDZ domain scaffold protein that exhibits GTPase activating protein binding activity

Expression: Body wall musculature; nerve ring; neurons and vulval muscle

Previously reported phenotypes: *mpz-1* RNAi reduces 5-HT stimulated egg laying[19]

#### Human Orthologs:

*MPDZ*

Associated disease(s) :
Hydrocephalus, nonsyndromic, 2, 615219 (AR)

#### Results

4957/8289 significant features vs ($p < 0.05$, block permutation t-test, 10000 permutations)

Key phenotype(s): Shorter; decreased curvature of the head/neck but increased curvature of hips/midbody; increased speed (hypersensitive to blue light); hyperactive; sustained forward escape response; increased rate of reversals (fraction of backwards moving worms)

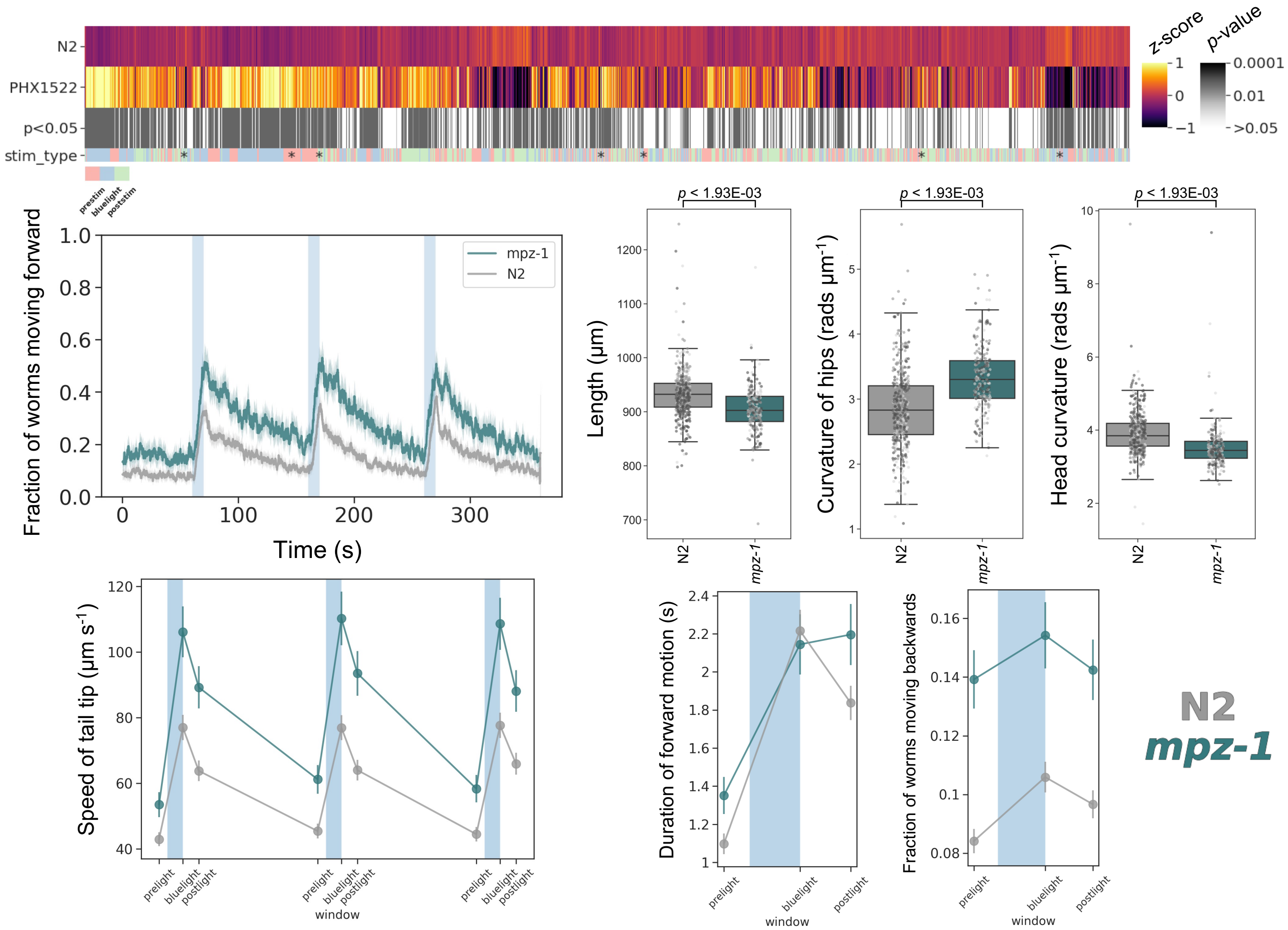

### *nca-2(syb1612)*

Strain name: PHX1612
Wild-type gene length: 8377bp
Mutation: 6003bp deletion (first 13.5 exons)

#### *C. elegans* Description

Function: Cation leak channel activity; interacts with *unc-77* and *unc-80*

Expression: Intestine; pharyngeal neurons and somatic nervous system

Previously reported phenotypes: Locomotion deficits; fainter in liquid; sensitive to halothane and isoflurane[15]

#### Human Orthologs:

*NALCN*

Associated disease(s):
Congenital contractures of the limbs and face, hypotonia, and developmental delay, 616266 (AD); Hypotonia, infantile, with psychomotor retardation and characteristic facies 1, 615419 (AR)

#### Results

2522/8289 significant features vs N2 ($p < 0.05$, block permutation t-test, 10000 permutations)

Key phenotype(s): Smaller; decreased speed; increased curvature/angular velocity of head only; no difference in baseline locomotion but an increased, short-lived, reversal response to blue light; increased post-stimulation pausing, “fainting” phenotype

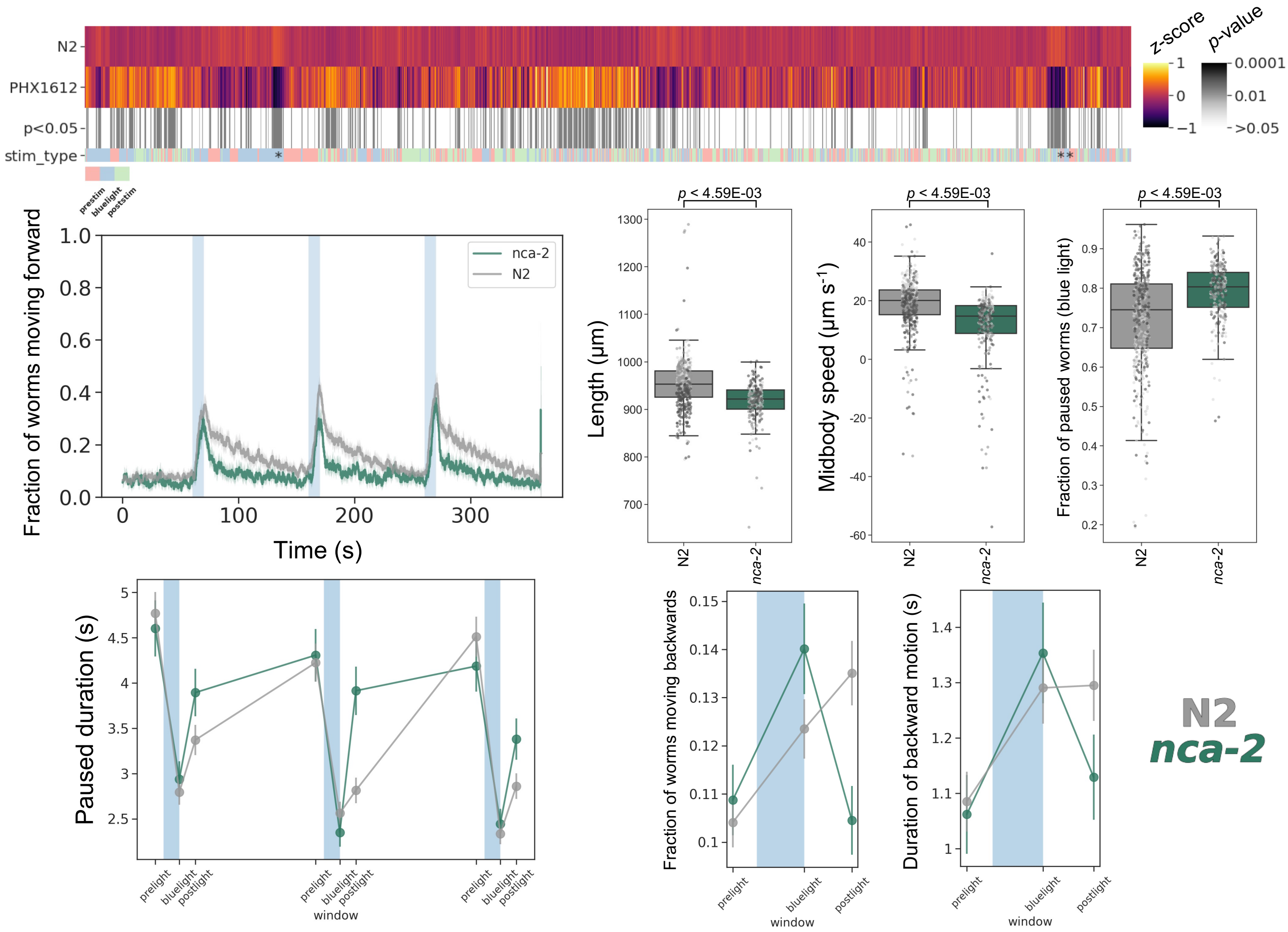

### *pink-1(syb1546)*

Strain name: PHX1546
Wild-type gene length: 1507bp
Mutation: 1381bp deletion (second half exon 2 to exon 6)

#### *C. elegans* Description

<u>Function</u>: Serine/threonine-specific protein kinase

<u>Expression</u>: Hermaphrodite distal tip cell; intestine; neurons; pharynx and vulva

<u>Previously reported phenotypes</u>: Required for: normal response to oxidative stress; mitochondrial homeostasis; correct CAN neurite outgrowth and wild type brood sizes. Mutant phenotypes are suppressed by mutation in *lrk-1* (LRRK2 ortholog)[20-22]

#### Human Orthologs:

*PINK1*

<u>Associated disease(s):</u>
Parkinson disease 6, early onset, 605909 (AR)

#### Results

0/8289 significant features vs N2 ($p < 0.05$, block permutation t-test, 10000 permutations)

Key phenotype(s): No significant phenotype detected

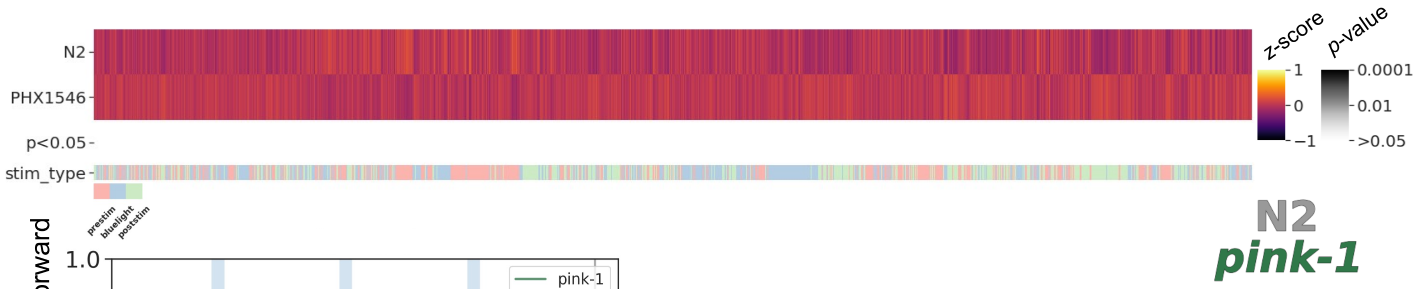


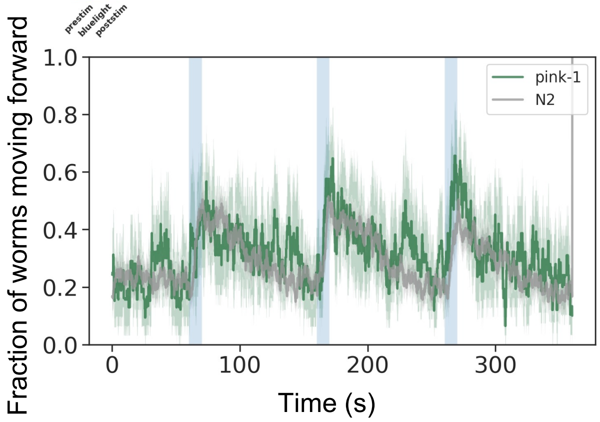

### ***snf-11(syb2683)***

Strain name: PHX2683
Wild-type gene length: 2283bp
Mutation: 1543bp deletion (exon 1 to second half of exon 4)

#### *C. elegans* Description

Function: Sodium/chloride ion-coupled high affinity GABA transporter; involved in gamma-aminobutyric acid transport

Expression: GLR; enteric muscle; head mesodermal cell; neurons and ventral nerve cord

Previously reported phenotypes: Abnormal head foraging and sinusoidal movements[15]

#### Human Orthologs:

*SLC6A1*

Associated disease(s):
Myoclonic-atonic epilepsy, 616421 (AD)

#### Results

3601/8289 significant features vs N2 ($p$ < 0.05, block permutation t-test, 10000 permutations)

Key phenotype(s): slightly less active; low frequency head bends (decreased time derivative of curvature); increased speed of head; increased foraging behaviour (time derivative of radial velocity); strong, but short-lived, backwards escape response to blue light stimulation

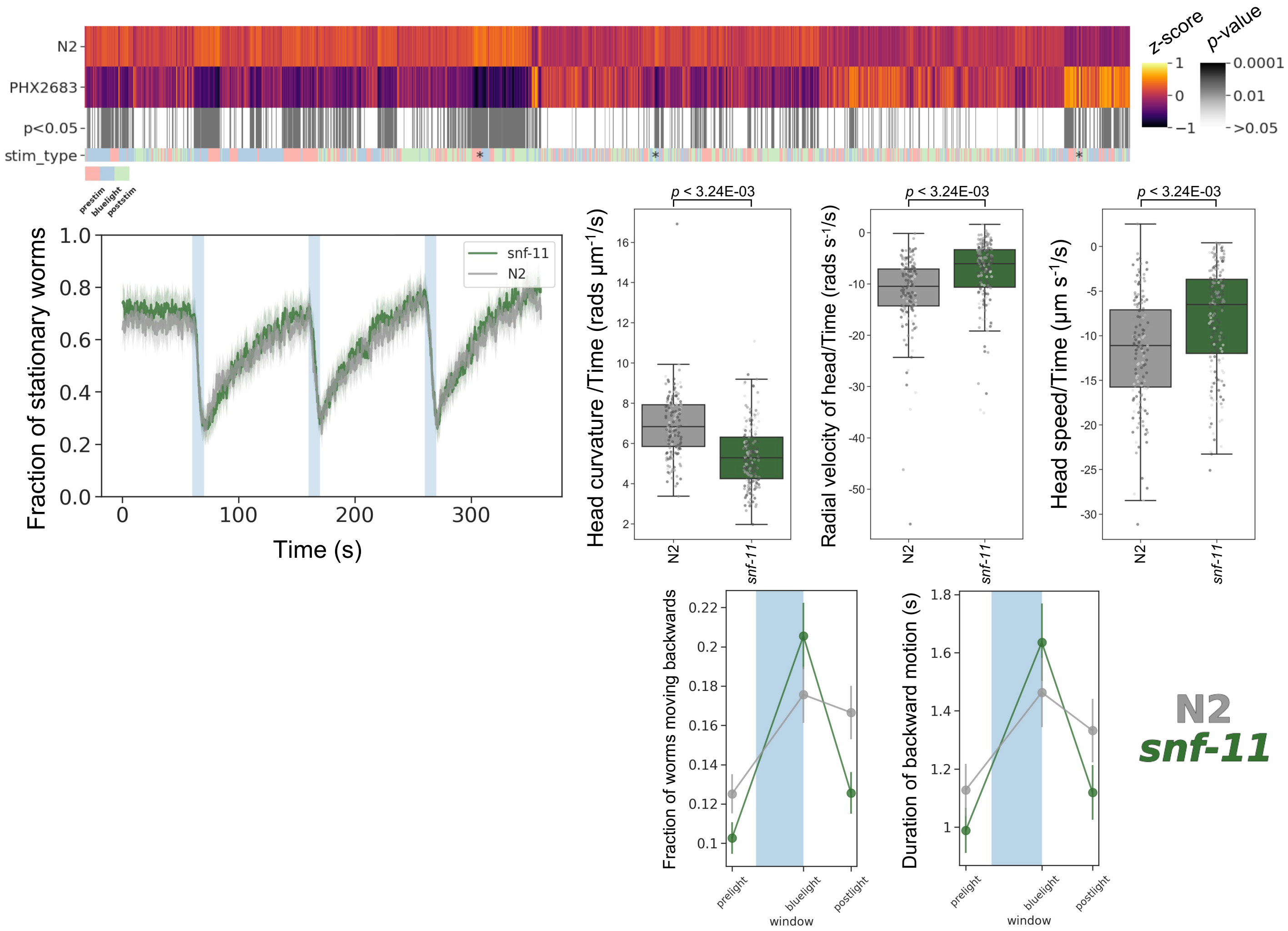

### ***snn-1(syb2695)***

Strain name: PHX2695
Wild-type gene length: 2682bp
Mutation: 1835bp deletion (2 exons)

#### *C. elegans* Description

Function: Synapsin, involved in neuromuscular synaptic transmission

Expression: All neurons; localises to presynaptic regions

Previously reported phenotypes: Aldicarb resistance; defective synaptic vesicle clustering[13,23]

#### Human Orthologs:

*SYN1; SYN2*

Associated disease(s):
Epilepsy, X-linked recessive and dominant (XLR, XLD), with variable learning disabilities and behaviour disorders, 300491; Schizophrenia susceptibility, 181500 (AD)

#### Results

1633/8289 significant features vs N2 (*p* < 0.05, block permutation t-test, 10000 permutations)

Key phenotype(s): Less active; shallow head movements (decreased time derivative of curvature); increased head speed; increased foraging behaviour (time derivative of radial velocity); increased reversal response to blue light that diminishes upon repeated exposure

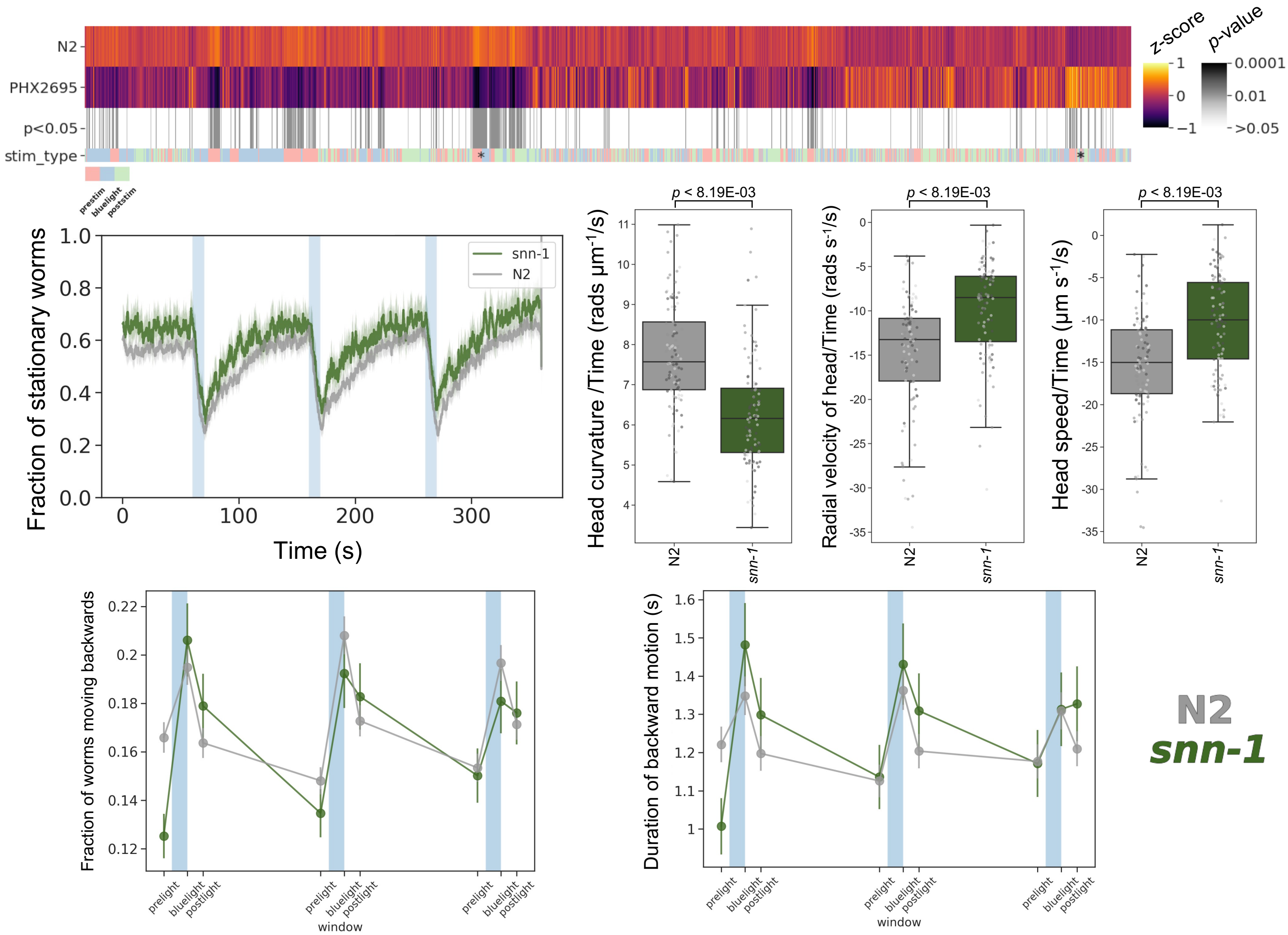

### ***tmem-231(syb1575)***

Strain name: PHX1575
Wild-type gene length: 2033bp
Mutation: 1253bp deletion (exon 1 to beginning of exon 7)

#### *C. elegans* Description

<u>Function</u>: Involved in non-motile cilium assembly

<u>Expression</u>: Amphid neurons; ciliated neurons; inner labial neurons; outer labial neurons and phasmid neurons

<u>Previously reported phenotypes</u>: No previous phenotype reported

#### Human Orthologs:

*TMEM231*

<u>Associated disease(s):</u>
Joubert syndrome 20, 614970 (AR); Meckel syndrome 11, 615397 (AR)

#### Results

811/8289 significant features vs N2 ($p$ < 0.05, block permutation t-test, 10000 permutations)

Key phenotype(s): Slightly less active during peristimulus recording; increased head curvature and angular velocity of tail (during peristimulus only); increased speed upon stimulation with blue light; sustained backwards escape response to blue light

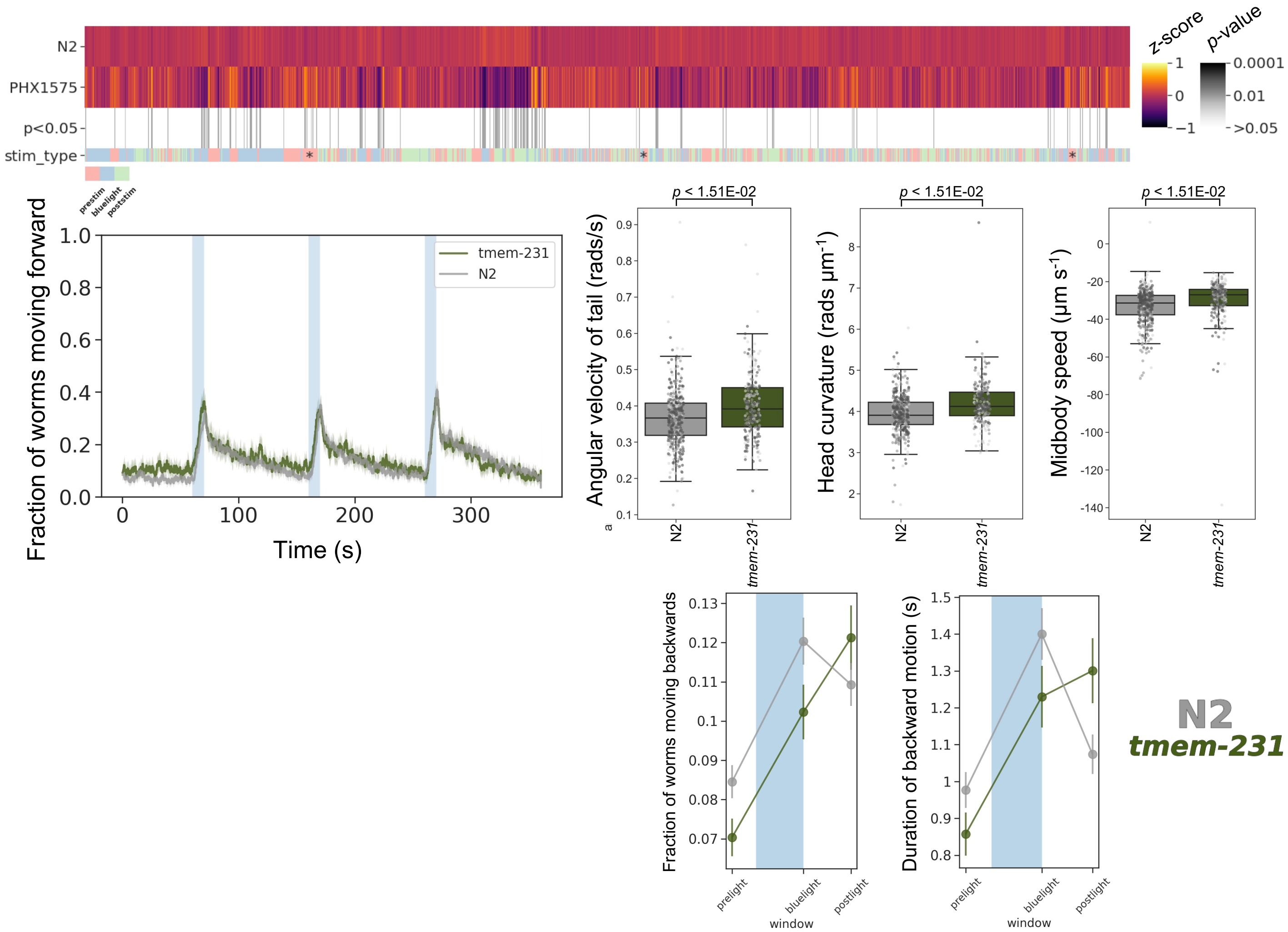

### *tub-1(syb1563)*

Strain name: PHX1563
Wild-type gene length: 1964bp
Mutation: 1319bp deletion (exon 1 to beginning of exon 4)

#### *C. elegans* Description

<u>Function</u>: Encodes TUBBY homolog and interacts with a 3-ketoacyl-coA thiolase, affecting lipid accumulation; involved in localisation of G-protein-coupled receptors in nonmotile cilia

<u>Expression</u>: Cytoplasm of ciliated cells

<u>Previously reported phenotypes</u>: Extended lifespan; increased fat content[24-25]

#### Human Orthologs:

*TUB, TULP1*

<u>Associated disease(s):</u>
Retinal dystrophy and obesity, 616188 (AR); Leber congenital amaurosis 15, 613843 (AR); Retinitis pigmentosa 14, 600132 (AR)

#### Results

4567/8289 significant features vs N2 ($p < 0.05$, block permutation t-test, 10000 permutations)

Key phenotype(s): Shorter and fatter; decreased curvature; deeper body bends (increased time derivative of curvature); hyperactive prior to simulation; decreased speed; attenuated, but sustained, sustained forward escape response; little difference in backwards motion/reversals

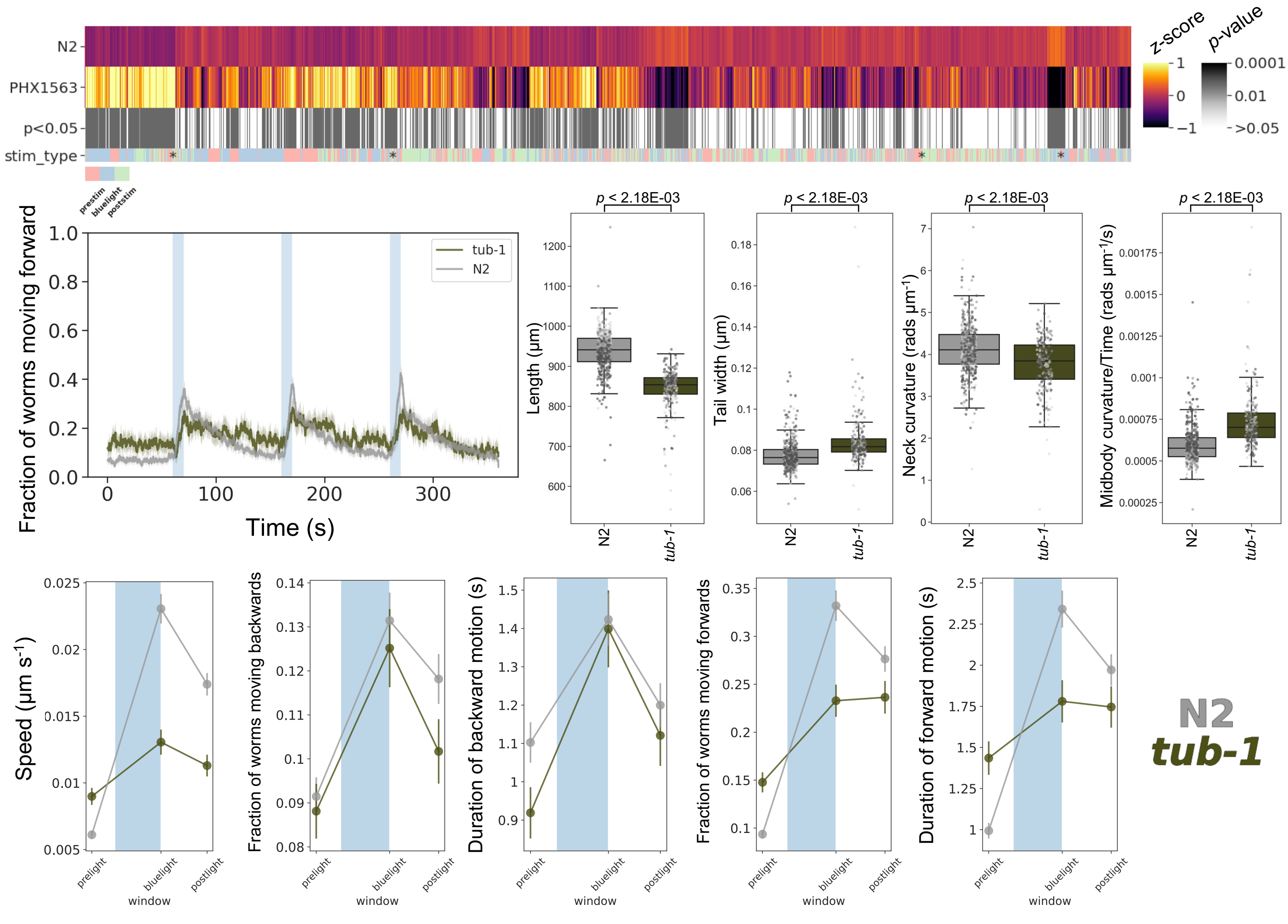

### ***unc-25(syb1651)***

Strain name: PHX1651
Wild-type gene length: 1706bp
Mutation: 1304bp deletion (exons 2-4)

#### *C. elegans* Description

<u>Function</u>: Encodes glutamic acid decarboxylase (inolved in GABA biosynthesis)

<u>Expression</u>: EF neuron and neurons

<u>Previously reported phenotypes</u>: GABA deficient; shrinker; defective defecation; irregular head thrashes; sensitive to PTZ, fluoxetine and aldicarb[26-32]

#### Human Orthologs:

*GAD1*

<u>Associated diseases(s):</u>
Cerebral palsy, spastic quadriplegic, 1, 603513 (AR)

#### Results

1956/8289 significant features vs N2 ($p$ < 0.05, block permutation t-test, 10000 permutations)

Key phenotype(s): Developmentally delayed (2 days); shorter; deeper sinusoidal bends (increased time derivative of curvature); less active and slower; delayed, but sustained forward escape response to blue light exposure

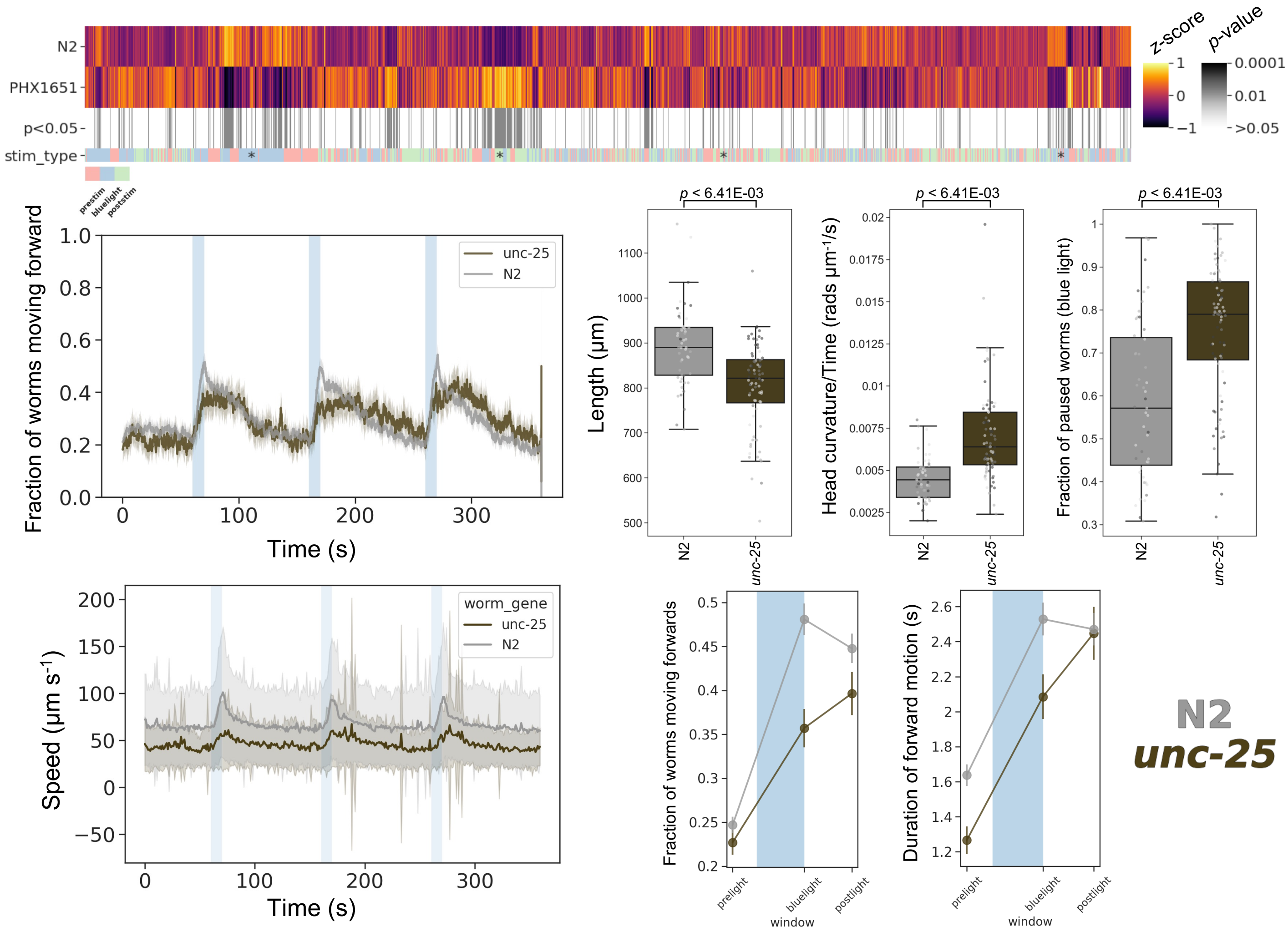

### *unc-43(syb2654)*

Strain name: PHX2654
Wild-type gene length: 1823bp
Mutation: 1246bp deletion (exon 1 to beginning of exon 3)

#### *C. elegans* Description

<u>Function</u>: Encodes CamKII-kinase; UNC-43 also interacts with the EGL-2 (ether-a-go-go potassium channel) to regulate it's activity

<u>Expression</u>: Neurons; oocytes; gonadal sheath cells; copulatory spicule

<u>Previously reported phenotypes</u>: Poor backing; slow moving; shrinker; defective egg laying; body wall muscle variant; sensitive to PTZ and aldicarb[30,33-36]

#### Human Orthologs:

*CAMK2A; CAMK2B*

<u>Associated disease(s):</u>
Intellectual disability, 53, 617798 (AD); Intellectual disability, 54, 617799 (AD)

#### Results

6858/8289 significant features vs N2 ($p < 0.05$, block permutation t-test, 10000 permutations)

Key phenotype(s): Small; increased curvature and head foraging while moving (time derivative of angular velocity); hyperactive; increased speed; increased and sustained escape response to blue light stimulation

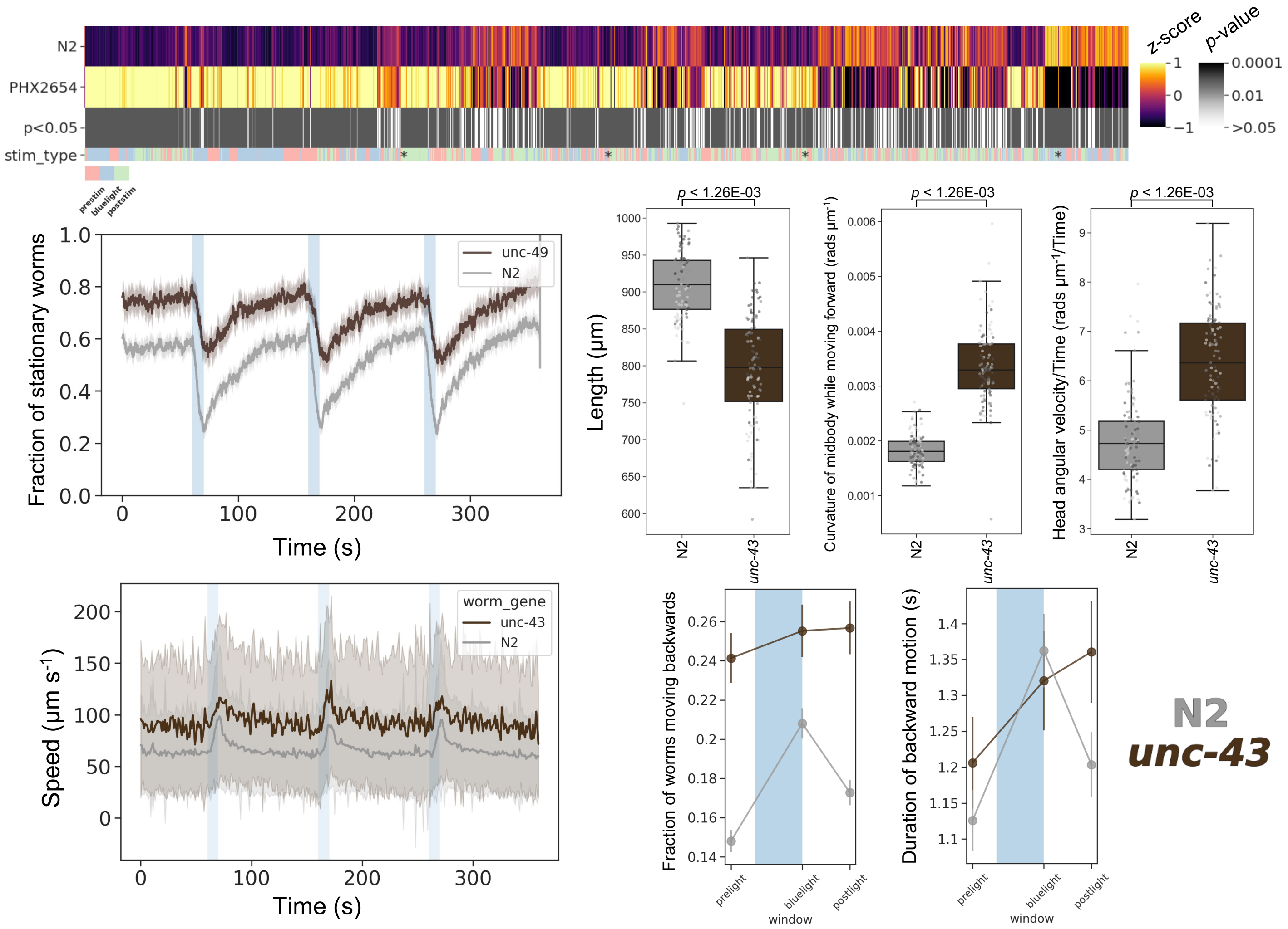

### *unc-49(syb2856)*

Strain name: PHX2856
Wild-type gene length: 8713bp
Mutation: 7765bp deletion (9 exons)

#### *C. elegans* Description

Function: Encodes subunits of heteromeric GABA receptor; UNC-49 is also required for postsynaptic GABA responsiveness

Expression: Expressed in anal sphincter muscle; body wall musculature; dorsal nerve cord; nerve ring and ventral nerve cord

Previously reported phenotypes: Decreased backing and thrashing; asymmetric body bends; small; shrinker; hypersensitive to fluoxetine and PTZ; resistant to muscimol[12,17-18,28,30-32,37-38]

#### Human Orthologs:

*GABRB1; GABRB2; GABRB3*

Associated disease(s):
Epileptic encephalopathy, early infantile, 43 and 45; Epileptic encephalopathy, infantile or early childhood, 2; Childhood absence epilepsy; Early onset epileptic encephalopathy

#### Results

4799/8289 significant features vs N2 (*p* < 0.05, block permutation t-test, 10000 permutations)

Key phenotype(s): Less active; smaller; decreased angular velocity of neck; increased curvature of midbody; decreased speed; sustained forward escape response, but attenuated backward escape response to blue light

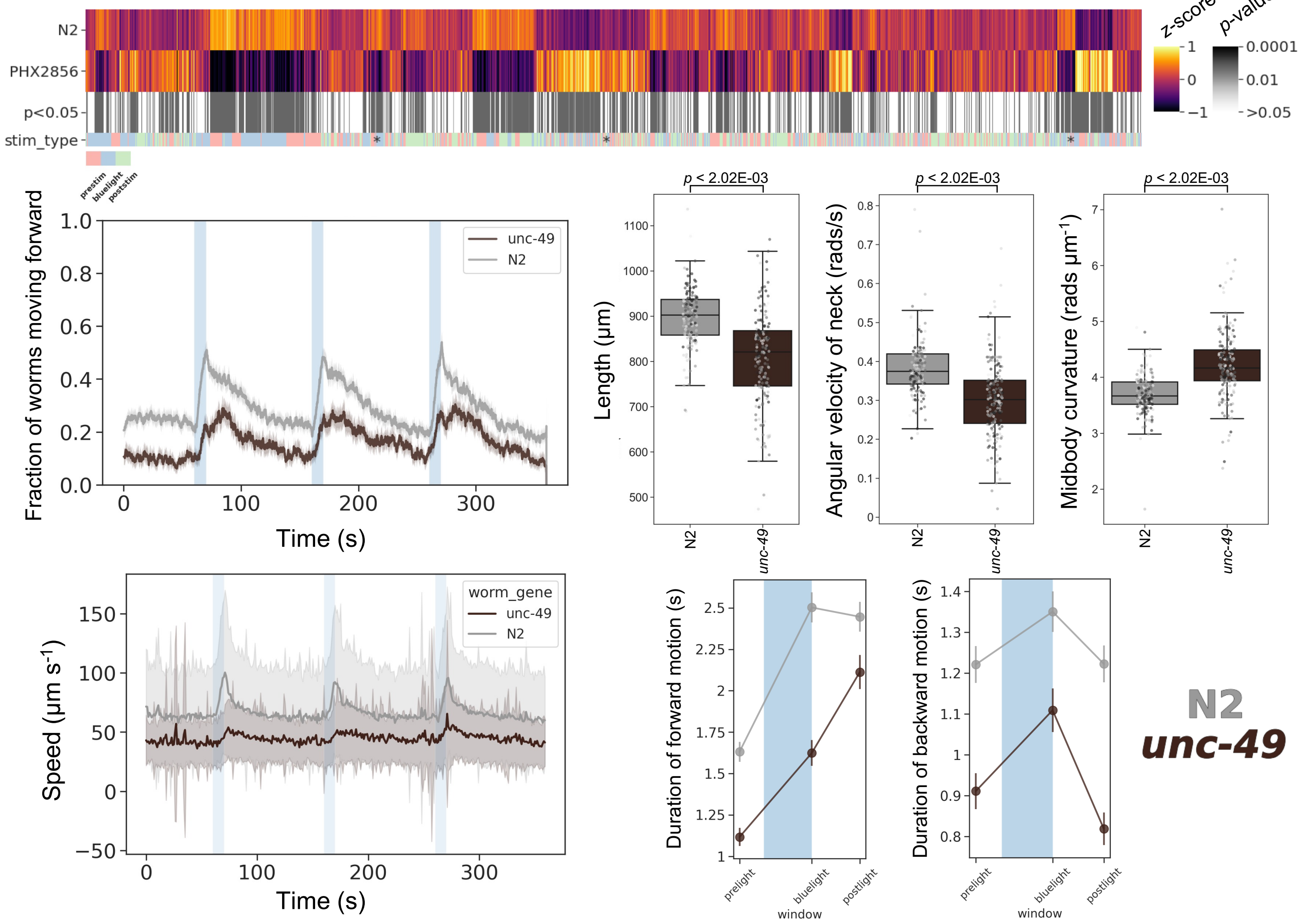

### *unc-77(syb1688)*

Strain name: PHX1688
Wild-type gene length: 2046bp
Mutation: 1407bp deletion (first 4 exons of NM171353.4)

#### *C. elegans* Description

<u>Function</u>: Cation-leak channel; interacts with *unc-80*, *unc-79* and *nca-2*

<u>Expression</u>: Cholinergic neurons; nerve ring; pharyngeal neurons and tail ganglion

<u>Previously reported phenotypes</u>: Decreased backing and increased sinusoidal movement[15]

#### Human Orthologs:

*NALCN*

<u>Associated disease(s):</u>
Congenital contractures of the limbs and face, hypotonia, and developmental delay, 616266 (AD); Hypotonia, infantile, with psychomotor retardation and characteristic facies 1, 615419 (AR)

#### Results

1011/8289 significant features vs N2 ($p < 0.05$, block permutation t-test, 10000 permutations)

Key phenotype(s): Decreased curvature of head/neck; stronger increase in speed in response to blue light; defective backing response, characterised by: decreased rate/duration of reversals, low frequency back bends (time derivative of curvature) and increase in time derivative of angular velocity related features across the body

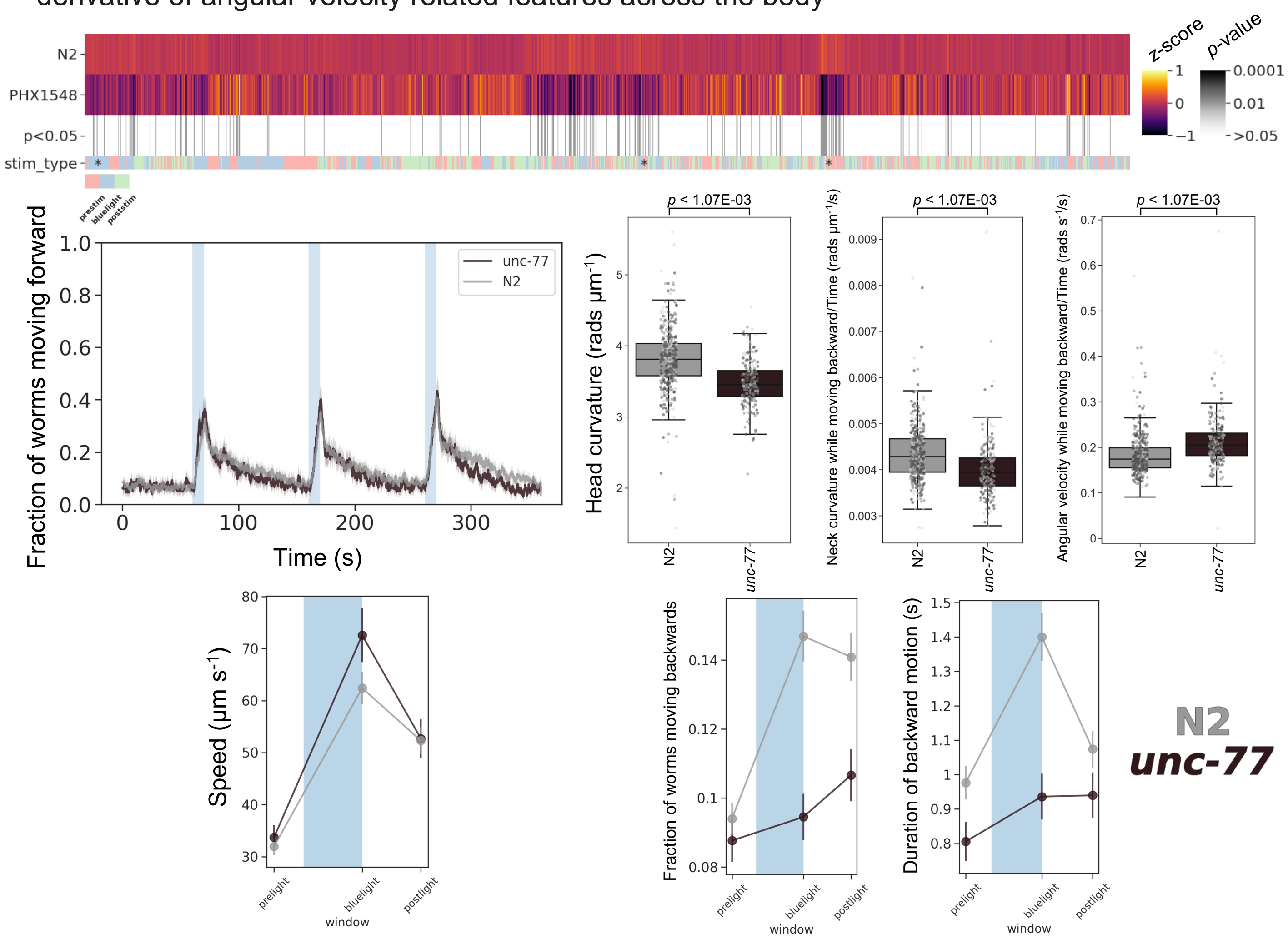

### *unc-80(syb1531)*

Strain name: PHX1531
Wild-type gene length: 1806bp
Mutation: 1367bp deletion (second half of exon 1 to middle of exon 4)

#### *C. elegans* Description

<u>Function</u>: Cation-leak channel; interacts with *unc-77*, *unc-79* and *nca-2*

<u>Expression</u>: Cholinergic neurons; excretory canal; hermaphrodite distal tip cell; non-striated muscle and somatic nervous system

<u>Previously reported phenotypes</u>: Sinusoidal, coiling and locomotion phenotypes; fainting in liquid; sensitivity to halothane, isoflurane and ethanol[15,39-40]

#### Human Orthologs:

*UNC80*

<u>Associated disease(s):</u>
Hypotonia, infantile, with psychomotor retardation and characteristic facies 2, 616801 (AR)

#### Results

4868/8289 significant features vs N2 ($p < 0.05$, block permutation t-test, 10000 permutations)

Key phenotype(s): Shorter; decreased curvature and speed; less active; attenuated blue light response (e.g., lack of backward escape response); and fainting (increased pausing) following cessation of blue light stimulation

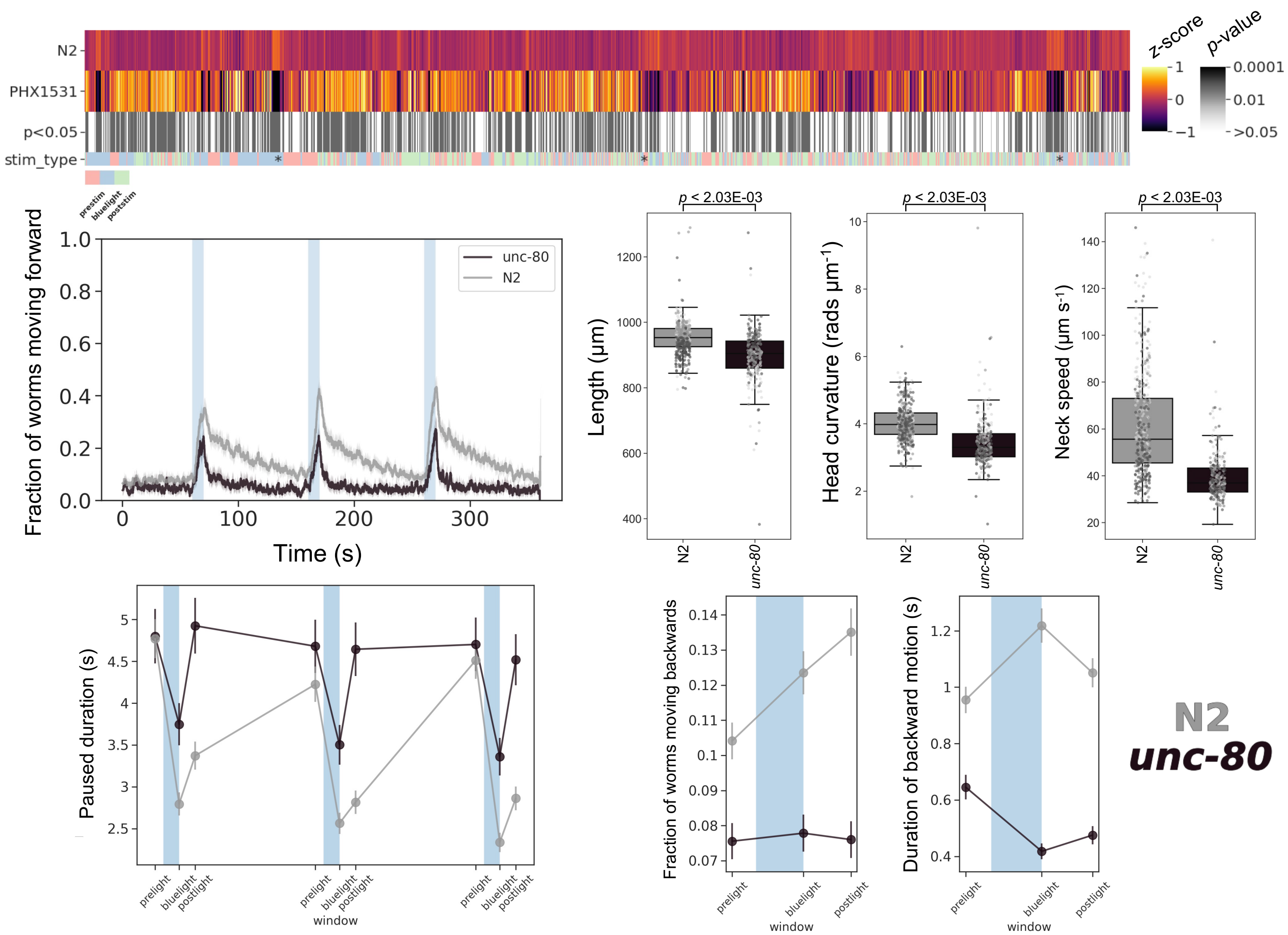
