## Supplementary Figure 1 for "Systematic creation and phenotyping of Mendelian disease models in *C. elegans*: towards large-scale drug repurposing"

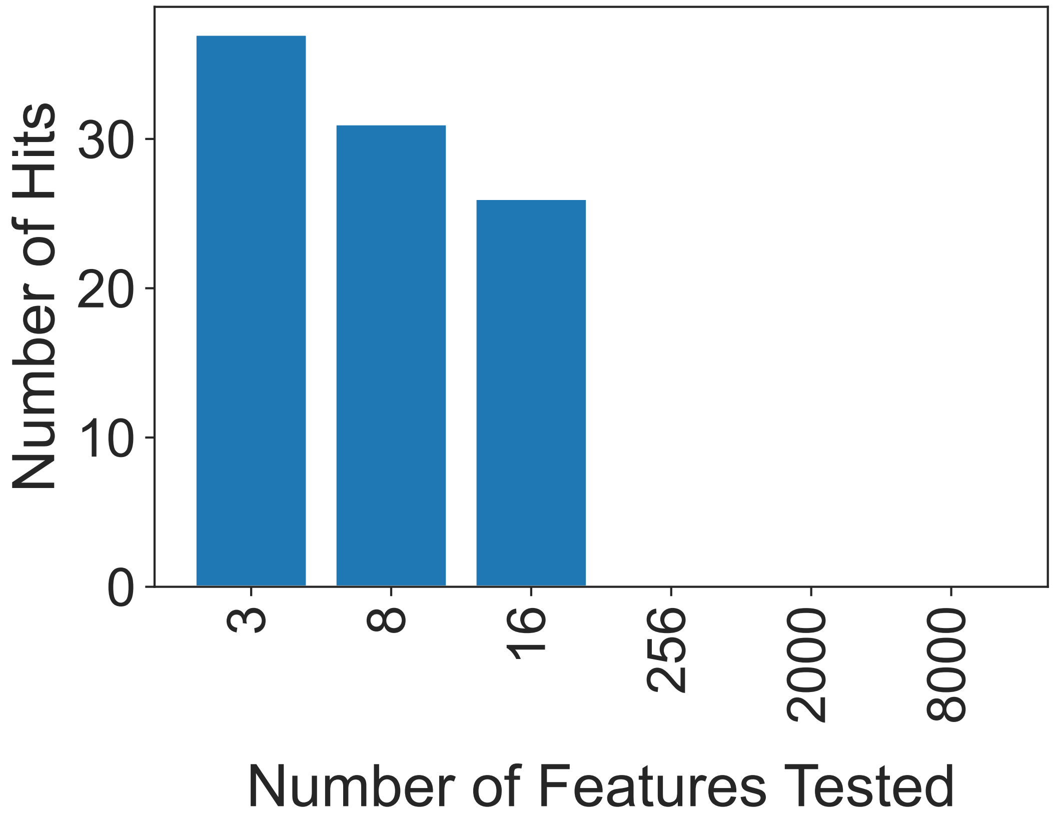


**Supplementary Figure 1: Number of initial compound hits detected when analysing increasing numbers of features.** The number of compound hits (as defined in the main text of this study) identified when performing statistical analysis (as detailed in the methods section) across the hand-selected core behavioural features (3), the entire feature set (8000) and for pre-defined Tierpsy feature sets containing an increasing number of features (Tierpsy-8, Tierpsy-16, Tierpsy-256 and Tierpsy-2k). There is a complete loss of statistical power when analysing >256 features using a low number of replicates ($n < 9$) and no statistically significant differences can be identified between the different drug treatments and the mutant or wild-type control after correcting for multiple comparisons.
