## Supplementary Figure 2 for "Systematic creation and phenotyping of Mendelian disease models in *C. elegans*: towards large-scale drug repurposing"

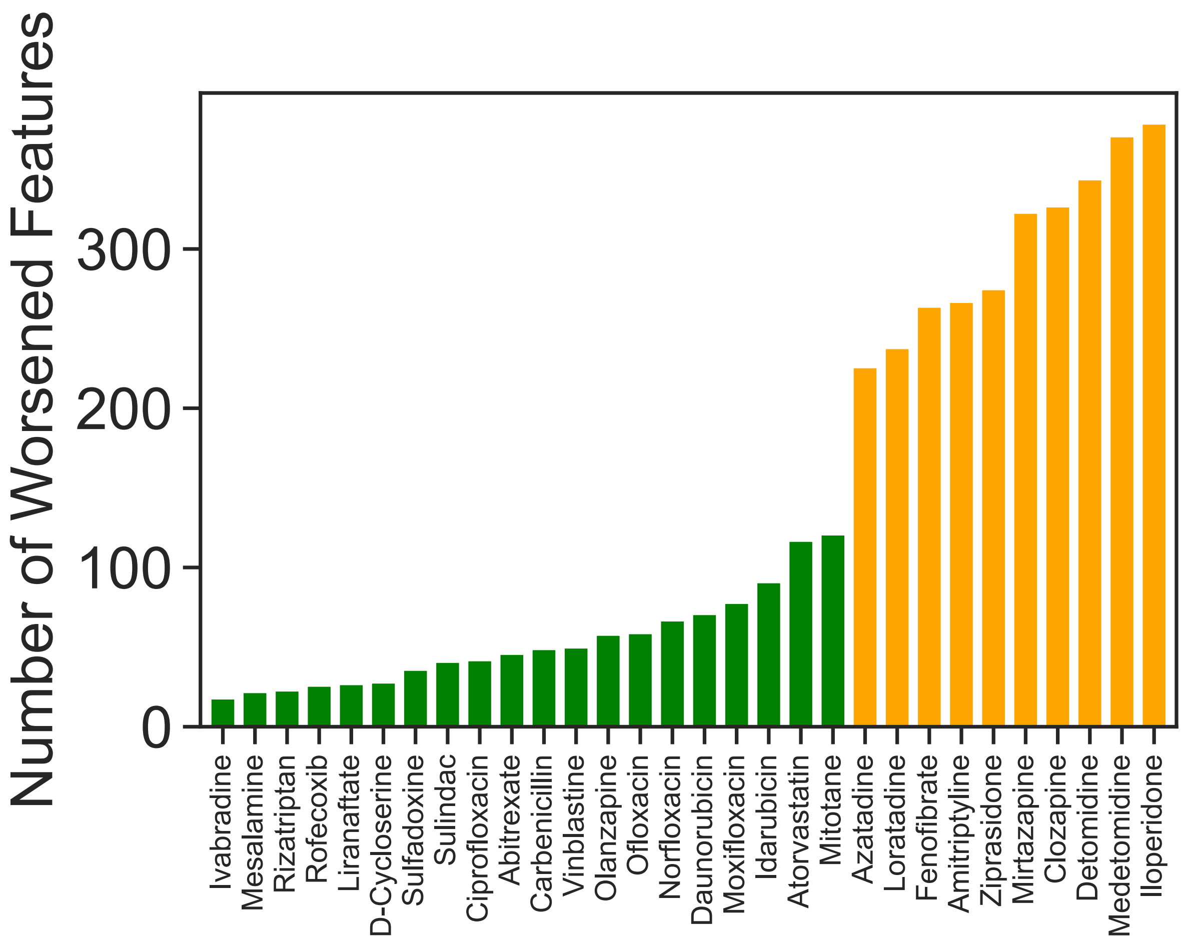


**Supplementary Figure 2: Behavioural side effects across a reduced behavioural featureset**
Total number of behavioural 'side effects' within a reduced set of features extracted by Tierpsy, Tierpsy-256, following treatment of *unc-80*(*syb1531*) with the 30 compounds in the confirmation screen. Side effects are defined as features that are not significant between *unc-80* mutants and wild-type N2 worms treated with 1% DMSO but where there is a significant difference between *unc-80* mutants treated with a drug compared to N2. The plot is coloured as described in Figure 5.
