## Supplementary Table 1 for "Systematic creation and phenotyping of Mendelian disease models in *C. elegans*: towards large-scale drug repurposing"

| Worm Gene | Databases Predicting Gene is a Human Ortholog | Ensembl ID | WormBase ID | No. Programs Predicting Orthology | Genetic similarity (%) | Blast E-value |
|---|---|---|---|---|---|---|
| *bbs-1* | Ensembl Compara 87-89\|Homologene\|InParanoid\|OrthoInspector | ENSG00000174483 | WBGene00000241 | 4 | 98.6 | 2.00E-77 |
| *snf-11* | Ensembl Compara 87-89\|Homologene\|InParanoid\|OrthoInspector\|OrthoMCL | ENSG00000157103 | WBGene00004910 | 5 | 98.2 | 0.00E+00 |
| *unc-25* | Ensembl Compara 87-89\|Homologene\|InParanoid\|OMA\|OrthoInspector\|OrthoMCL | ENSG00000128683 | WBGene00006762 | 6 | 96.9 | 0.00E+00 |
| *tmem-231* | Ensembl Compara 87-89\|InParanoid\|OMA\|OrthoInspector\|OrthoMCL | ENSG00000205084 | WBGene00020825 | 5 | 95.7 | 3.00E-12 |
| *unc-49* | OrthoInspector | ENSG00000101958 | WBGene00006784 | 1 | 94.1 | 1.00E-98 |
| *nca-2* | Ensembl Compara 87-89\|Homologene\|InParanoid\|OMA\|OrthoInspector\|OrthoMCL | ENSG00000102452 | WBGene00003558 | 6 | 93.9 | 0.00E+00 |
| *gpb-2* | Ensembl Compara 87-89\|Homologene\|InParanoid\|OMA\|OrthoInspector\|OrthoMCL | ENSG00000069966 | WBGene00001680 | 6 | 93 | 5.00E-172 |
| *unc-77* | Ensembl Compara 87-89\|Homologene\|InParanoid\|OrthoInspector\|OrthoMCL | ENSG00000102452 | WBGene00006809 | 5 | 91.9 | 0.00E+00 |
| *kcc-2* | Ensembl Compara 87-89\|InParanoid\|OrthoInspector\|OrthoMCL | ENSG00000124140, ENSG00000140199 | WBGene00019205 | 4 | 90.5 | 0.00E+00 |
| *glc-2* | Ensembl Compara 87-89\|OrthoInspector\|OrthoMCL | ENSG00000145888, ENSG00000109738 | WBGene00001592 | 3 | 90.1 | 2.00E-91 |
| *avr-14* | Ensembl Compara 87-89\|Homologene\|OMA\|OrthoInspector\|OrthoMCL | ENSG00000145888, ENSG00000109738 | WBGene00000232 | 5 | 90 | 9.00E-103 |
| *bbs-2* | Ensembl Compara 87-89\|Homologene\|InParanoid\|OMA\|OrthoInspector\|OrthoMCL | ENSG00000125124 | WBGene00000242 | 6 | 89.4 | 4.00E-119 |
| *unc-43* | Ensembl Compara 87-89\|InParanoid\|OrthoInspector | ENSG00000070808, ENSG00000058404 | WBGene00006779 | 3 | 83 | 0.00E+00 |
| *cat-4* | Ensembl Compara 87-89\|Homologene\|InParanoid\|OMA\|OrthoInspector\|OrthoMCL | ENSG00000131979 | WBGene00000298 | 6 | 82.5 | 2.00E-98 |
| *add-1* | Ensembl Compara 87-89\|InParanoid\|OrthoInspector\|OrthoMCL | ENSG00000148700, ENSG00000087274 | WBGene00000072 | 4 | 79.9 | 2.00E-119 |
| *glr-4* | Ensembl Compara 87-89\|Homologene\|OrthoInspector\|OrthoMCL | ENSG00000164418 | WBGene00001615 | 4 | 79.1 | 6.00E-151 |
| *glr-1* | Ensembl Compara 87-89\|Homologene\|InParanoid\|OrthoInspector\|OrthoMCL | ENSG00000125675, ENSG00000164418 | WBGene00001612 | 5 | 78 | 4.00E-180 |
| *figo-1* | Ensembl Compara 87-89\|InParanoid\|OMA\|OrthoInspector\|OrthoMCL | ENSG00000112367 | WBGene00007912 | 5 | 74.4 | 8.00E-160 |
| *pink-1* | Ensembl Compara 87-89\|Homologene\|InParanoid\|OrthoInspector\|OrthoMCL | ENSG00000158828 | WBGene00017137 | 5 | 72.1 | 9.00E-57 |
| *snn-1* | Ensembl Compara 87-89\|InParanoid\|OrthoInspector\|OrthoMCL | ENSG00000008056, ENSG00000157152 | WBGene00004913 | 4 | 66.2 | 1.00E-43 |
| *cat-2* | Ensembl Compara 87-89\|Homologene\|OMA\|OrthoMCL | ENSG00000180176 | WBGene00000296 | 4 | 62.6 | 8.00E-126 |
| *tub-1* | Ensembl Compara 87-89\|Homologene\|InParanoid\|OrthoInspector\|OrthoMCL | ENSG00000166402, ENSG00000112041 | WBGene00006655 | 5 | 61.3 | 3.00E-105 |
| *unc-80* | Ensembl Compara 87-89\|InParanoid\|OMA\|OrthoInspector\|OrthoMCL | ENSG00000144406 | WBGene00006812 | 5 | 42.8 | 0.00E+00 |
| *dys-1* | InParanoid\|OrthoInspector\|OrthoMCL | ENSG00000198947 | WBGene00001131 | 3 | 30.9 | 4.00E-169 |
| *mpz-1* | Ensembl Compara 87-89\|InParanoid\|OMA\|OrthoInspector\|OrthoMCL | ENSG00000107186 | WBGene00003404 | 5 | 16.1 | 3.00E-72 |

**Supplementary Table 1. Orthology program predictions and genetic similarities of the panel of disease model mutants.**
Table indicating the unique Wormbase and Ensembl database ascension numbers for every *C. elegans* gene in our panel of disease model mutants. Alongside denoting which (and how many) gene orthology databases predict that the worm gene is an orthologous to a human gene, and the genetic similarity and Blast-E score for each *C. elegans* gene.
